# Premature aging in aneuploid yeast is caused in part by aneuploidy-induced defects in Ribosome Quality Control

**DOI:** 10.1101/2024.06.22.600216

**Authors:** Leah E. Escalante, James Hose, Hollis Howe, Norah Paulsen, Michael Place, Audrey P. Gasch

## Abstract

Premature aging is a hallmark of Down syndrome, caused by trisomy of human chromosome 21, but the reason is unclear and difficult to study in humans. We used an aneuploid model in wild yeast to show that chromosome amplification disrupts nutrient-induced cell-cycle arrest, quiescence entry, and healthy aging, across genetic backgrounds and amplified chromosomes. We discovered that these defects are due in part to aneuploidy-induced dysfunction in Ribosome Quality Control (RQC). Compared to euploids, aneuploids entering quiescence display aberrant ribosome profiles, accumulate RQC intermediates, and harbor an increased load of protein aggregates. Although they have normal proteasome capacity, aneuploids show signs of ubiquitin dysregulation, which impacts cyclin abundance to disrupt arrest. Remarkably, inducing ribosome stalling in euploids produces similar aberrations, while up-regulating limiting RQC subunits or proteins in ubiquitin metabolism alleviates many of the aneuploid defects. Our results provide implications for other aneuploidy disorders including Down syndrome.

## INTRODUCTION

Chromosome amplification, here referred to as aneuploidy, is very detrimental during mammalian development and a leading cause of infertility in humans, for reasons that remain incompletely understood. Trisomy of human chromosome 21 that causes Down syndrome (DS) is the only autosomal aneuploidy viable into adulthood, thanks to decades of medical advances to improve health^1,2^. However, one of the most penetrant hallmarks of DS and a remaining medical concern is premature aging, including premature skin wrinkling and hair loss, defects in tissue regeneration, and early-onset neurodegeneration including Alzheimer’s^3–6^. The reasons for premature systemic aging are largely a mystery, in part because the underlying cellular consequences of chromosome amplification remain unknown despite over 65 years of study.

Budding yeast *Saccharomyces cerevisiae* has been an excellent model to understand the cellular stress of chromosome amplification. Previous seminal work studying an aneuploidy-sensitized laboratory strain revealed that chromosome duplication in this strain produces myriad defects in cell metabolism, stress response, and management of protein homeostasis known as proteostasis^7–12^. In contrast, wild isolates of yeast studied to date are much more tolerant of aneuploidy, with milder growth defects during logarithmic growth and little sign of proteostasis stress unless cells are further taxed^13–15^. The reason for the phenotypic differences across strains is traced to RNA binding protein Ssd1, which is functional in wild strains but hypomorphic in the sensitized W303 lab strain^13,16^. Ssd1 is involved in translational control and mRNA localization, and binds to several hundred transcripts encoding diverse functions^17–21^. Deletion of *SSD1* from wild strains renders cells highly sensitive to chromosome amplification, with cells showing many of the signatures of the sensitized lab strain including proteostasis stress^13,22^. Recent modeling work from our lab points to a defect in translational regulation in *ssd1Δ* aneuploids^22,23^. Furthermore, aneuploid yeast, especially aneuploid strains lacking *SSD1*, are sensitive to translational inhibitors, including nourseothricin (NTC) that binds the ribosome and disrupts translation elongation^13,22,24^. Our hypothesis is that wild *S. cerevisiae* isolates can handle the stress of chromosome duplication, in part through Ssd1-dependent mechanisms, but cells may be close to their buffering capacity under standard growth conditions^13^.

While studying wild aneuploid strains, we made an important discovery: although these aneuploids proliferate with mild growth delay during exponential growth, they have a major defect entering and maintaining quiescence induced by nutrient depletion. Quiescence is an important state conserved across taxa, in which cells exit the cell cycle upon specific cues but retain the ability to re-initiate proliferation at a later time^25–28^. Quiescence is necessary for proper development and critical for growth control, tissue homeostasis, and cellular longevity^26,29–31^. In fact, people with DS and animal DS models have defects maintaining quiescent stem cells, a deficiency that may underlie other phenotypes associated with premature aging^32–34^. Haploid yeast has served as an important model for understanding the quiescent state and defining key stages of the process^28,35^.

Here we show that chromosome amplification in yeast disrupts quiescence and lifespan due in part to defects in the Ribosome Quality Control (RQC) pathway. This pathway detects, disassembles, and clears collided ribosomes and the incomplete nascent polypeptides associated with them^36–38^. Part of the clearance mechanism involves non-mRNA-templated addition of alanine and threonine residues called ‘CAT’ tails to the nascent-peptide C-terminus by Rqc2 (NEMF in humans), followed by ubiquitination by the E3 ligase Ltn1 (mammalian Listerin) that triggers proteasomal degradation^36,39–41^. Failure to clear stalled products, in particular CATylated peptides that are prone to aggregation, produces toxic aggregates and proteostasis collapse^41–43^. Defects in translation and RQC are known to accumulate with age^44–46^. Furthermore, RQC dysfunction contributes to neurodegeneration in mammals and human disease models, and neurons are particularly sensitive to protein aggregation and RQC defects^47–52^. In this work, we present a model for how aneuploidy produces translational errors and consequential protein aggregation, which disrupts several processes to accelerate aging.

## RESULTS

### Chromosomal duplication disrupts quiescence

In the course of ongoing investigations, we discovered that haploid derivatives of wild oak-soil strain YPS1009 with different chromosome duplications showed abnormal arrest and regrowth after nutrient exhaustion. To investigate systematically, we characterized each step along the progression to quiescence in a series of engineered YPS1009 aneuploid strains each with a full duplication of various chromosomes ^23^. In this strain background, euploid cells shift to respiration as glucose is depleted from the media (day 1 of culturing), arrest as unbudded cells in G0 (day 1-2), silence their transcriptome (day 1-7), and become small and dense (day 4-7). All of these steps are important for healthy lifespan^53–55^.

We found that all aneuploids tested had defects in these quiescence hallmarks, to varying degrees. One of the earliest steps is cell-cycle arrest upon nutrient exhaustion in a saturated culture. Whereas nearly 100% of euploid cells arrest as unbudded cells by 2 days of culturing, 1-12% of cells, depending on the amplified chromosome, showed morphology indicative of budding (**Fig 1A**), even though the cultures had completely exhausted glucose (**supplemental Fig S1A**). The arrest defect is not specific to YPS1009, as two other strain backgrounds with extra chromosomes showed similar defects (**Fig 1B**). In addition to the arrest defect, YPS1009 with an extra chromosome XIV (YPS1009_Chr14) showed an unusual morphology of bi-lobed cells with a single round nucleus in between, while YPS1009_Chr15 took on a multi-budded elongated state (**Fig 1C**). Importantly, YPS1009_Chr14 and _Chr15 aneuploids do not show these morphologies under standard conditions (**supplemental Fig S1B**), indicating that these morphologies are specific to nutrient starvation. Thus, chromosome duplication generally disrupts cell-cycle arrest upon nutrient exhaustion, albeit with some chromosome-specific effects (see Discussion).

**Figure 1.**
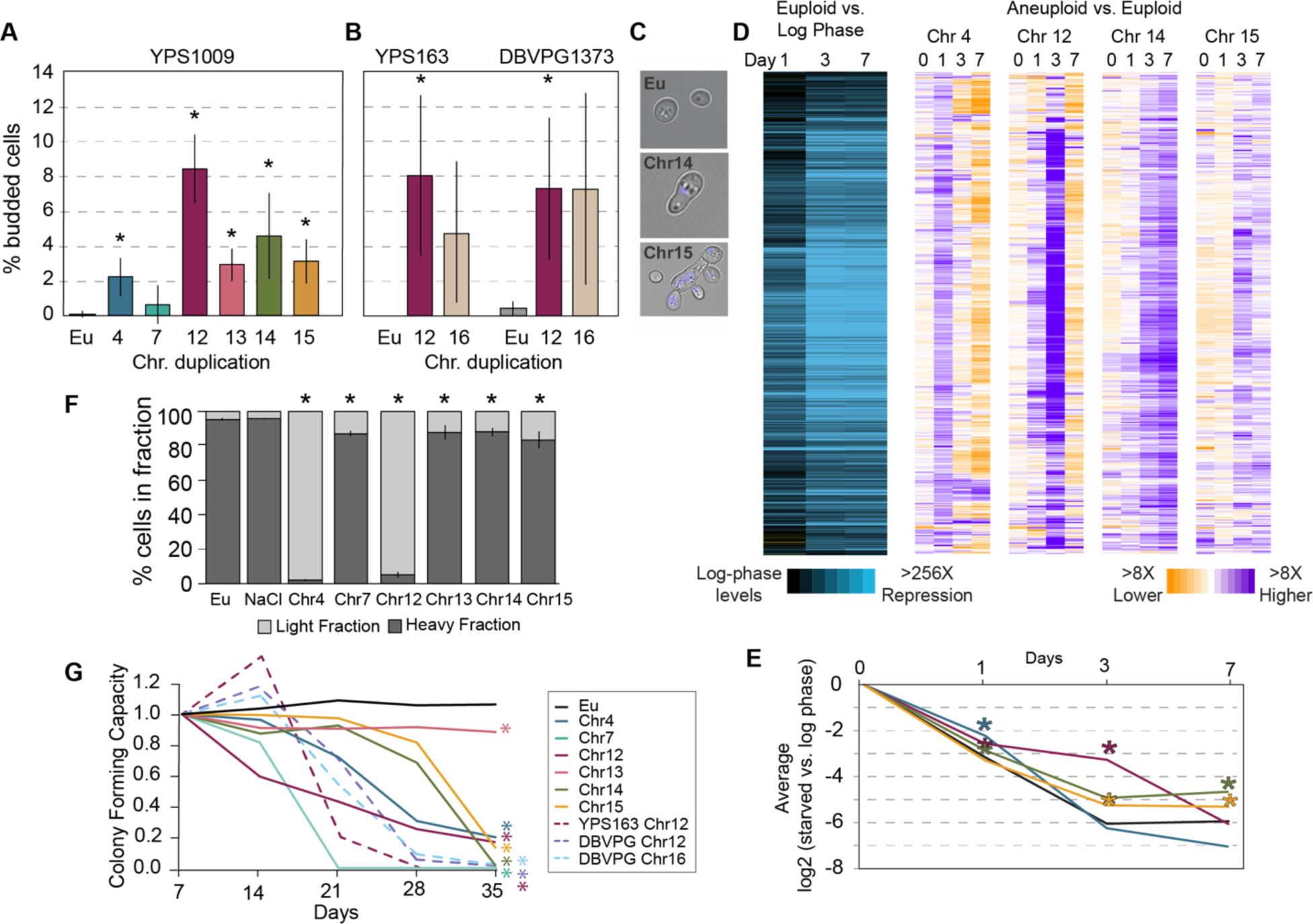
Chromosome duplication disrupts multiple signatures of quiescence. (A-B) Percent budding cells at 2 days in (A) YPS1009 (n>=3) or (B) oak strain YPS163 or vineyard strain DBVPG1373 (n=2-3) with chromosome duplications as indicated. Asterisk, p<0.05, T-test comparing aneuploids versus matched euploid (Eu). (C) Representative brightfield images from A. Blue represents DAPI staining. (D) Left: Replicate-averaged log_2_(fold change) expression of 1440 genes (rows) repressed in euploid but significantly higher in abundance (FDR < 0.05) in all four aneuploids compared to euploid, in at least one timepoint (see Methods). Right: The replicate-averaged log_2_(fold difference) in normalized transcript abundance for genes shown on the left, in each aneuploid versus euploid at that timepoint. Purple indicates a repression defect. (E) Average of all genes shown in D for strains in the key to the left. Asterisk, p < 1e-8, gene-paired T-test comparing each aneuploid to euploid. (F) Proportion of dense and light cells after 4 days (n=3-5). Asterisk, p<0.05, Chi-squared test. (G) Average fraction of colony forming units relative to day 7 (YPS1009 strains, n=3-6; YPS163 and DBVPG1373, n=2). Asterisk, p<0.05, paired T-test comparing aneuploids to euploid at day 35.

Aneuploid cultures also displayed defects in subsequent steps of quiescence, including the stereotypical transcriptome silencing associated with quiescence^54,56,57^. Euploid cultures repressed thousands of transcripts, >256X below log-phase levels, beginning 1 day past exponential phase and dropping to stable levels by day 3 (**Fig 1D-E**). We measured bulk transcriptomes of YPS1009 with a duplication of Chr4, Chr12, Chr14, or Chr15. The transcriptional profiles of all aneuploids were dysregulated to varying extents: we identified 1440 transcripts reduced in euploid cells but at significantly higher abundance in all four aneuploids, in at least one timepoint (FDR < 0.05, see Methods). Two of the aneuploids eventually reached euploid repression after a delay. Transcriptome silencing is known to be important for quiescence, since defective silencing shortens lifespan^58^. In addition to the silencing defect, aneuploidy also disrupted cell densification that occurs during this time frame. Unlike euploid cells that densify after 4 days of culturing, all aneuploids tested showed statistically significant effects, with YPS1009_Chr4 and _Chr12 showing the largest defect (**Fig 1F**).

Quiescence is fundamentally important for normal life span in yeast ^29,30,59,60^ – indeed, aneuploids have a substantially shorter chronological lifespan (**Fig 1G**). Euploids exhibited near 100% colony forming capacity over 5 weeks after nutrient exhaustion. Instead, all but one aneuploid tested exhibited a dramatically shorter lifespan, with few, if any, viable cells remaining by the end of the time course (**Fig 1G**). The lone exception, YPS1009_Chr13, showed lower colony forming capacity (60-70%) after only 1 day of culturing that remained stably lower over 5 weeks. The reduced lifespan was also seen in two other strain backgrounds, again indicating that the effect is not specific to YPS1009. Our results expand past studies showing that chromosome duplication in the aneuploidy-sensitized W303 lab strain shortens replicative life span^61^ and confirms that the effect is not due to loss of Ssd1, which is important for both quiescence and lifespan in euploid yeast^20,55,62,63^.

Collectively, our results show that chromosome amplification disrupts multiple signatures of quiescence and lifespan, independent of which chromosome is amplified but with some chromosome-specific effects. This strongly suggests a generalizable consequence of chromosome duplication on aging and lifespan, overlayed with chromosome-specific impacts (see Discussion).

### A genetic screen for lifespan extension implicates the RQC pathway

To understand the mechanisms of lifespan limitation, we conducted a genetic screen in YPS1009_Chr12 cells, using a low-copy, barcoded plasmid library expressing each of ∼5,000 yeast genes flanked with their native upstream and downstream sequences^64^. YPS1009_euploid or YPS1009_Chr12 cells transformed with the library were pooled separately, grown to saturation, and cultured for 28 days. Viable colonies were identified by plating cells after 1 and 28 days, and changes in relative plasmid abundance compared to the starting library were scored by barcode sequencing of the pools. Genes whose duplication is beneficial to viability will increase in relative abundance compared to the starting pool (FDR < 0.05, see Methods).

We identified 59 genes that reproducibly increased in the pool over 28 days in YPS1009_Chr12 (FDR < 0.05) and were at least 2-fold more enriched than in the euploid (see Methods). Twenty-two genes were enriched only at 28 days and not day 1 of culturing, implicating a specific impact on lifespan (Dataset 1). Nearly half of these genes encode proteins localized to the mitochondria, whose function is critical for quiescence and lifespan but known to be defective in aneuploid yeast and human cells^65–68^. The other 37 genes were enriched at both 28 days and 1 day of culturing, suggesting an early impact on culture growth and healthspan. Among these are genes already linked to lifespan from other studies, including sirtuin Hst2 tenuously linked to lifespan^69–71^, several genes involved in autophagy (*ATG12*, *POR1*), which is required for healthy aging^72,73^, and others discussed more below (*SGT1*, *SCP1)*^74,75^.

Among the most intriguing on the list of 37 genes was *RQC1*, a component of the RQC pathway that resolves stalled ribosomes^36,37,76^. This was interesting because aneuploid strains are sensitive to translational inhibitor nourseothricin (NTC) that binds the ribosome to disrupt translation^13,22,77–79^. Given that translation defects increase with cellular age^44–46^, these results raised the possibility that chromosome duplication induces defects in the RQC pathway.

### The RQC pathway is directly involved in aneuploidy-dependent quiescence defects

We explored the impact of specific perturbations to the RQC pathway. Both Rqc1 and Ltn1 are stoichiometrically limiting in yeast and mammals^36,80,81^. Remarkably, the defect in cell-cycle arrest was significantly alleviated simply by duplicating *RQC1* or *LTN1* on a plasmid, in nearly all aneuploids tested: significantly more cells in each culture arrested as unbudded cells by 2 days (**Fig 2A**). Previous work showed that, counterintuitively, deletion of *RQC2* can actually alleviate the stress of a dysfunctional RQC pathway, by preventing accumulation of toxic CATylated peptides that are prone to aggregation^43,51,82^. Indeed, deletion of *RQC2* partly alleviated the arrest defect in aneuploid strains. We also found that duplication of *HEL2*, encoding the RQC sensor of collided ribosomes^83,84^, alleviated the arrest defect. We were unable to test the impact of Hel2 deletion, since the YPS1009_Chr4 *HEL2* deletion strain was not viable and YPS1009_Chr12 *hel2Δ* cultures did not maintain aneuploidy; this suggests that these aneuploids are especially reliant on Hel2. We also confirmed that *RQC1* duplication improved cell densification of both YPS1009_Chr4 and _Chr12 aneuploids and significantly increased culture growth in terms of final optical density leading up to quiescence (**supplemental Fig S2A-B**). Mere duplication of these genes was not enough to extend lifespan (**supplemental Fig S2C**), but it did alleviate other signatures of aging and proteostasis stress (see more below). Thus, augmenting the RQC pathway can improve healthspan in aneuploid cells.

**Figure 2.**
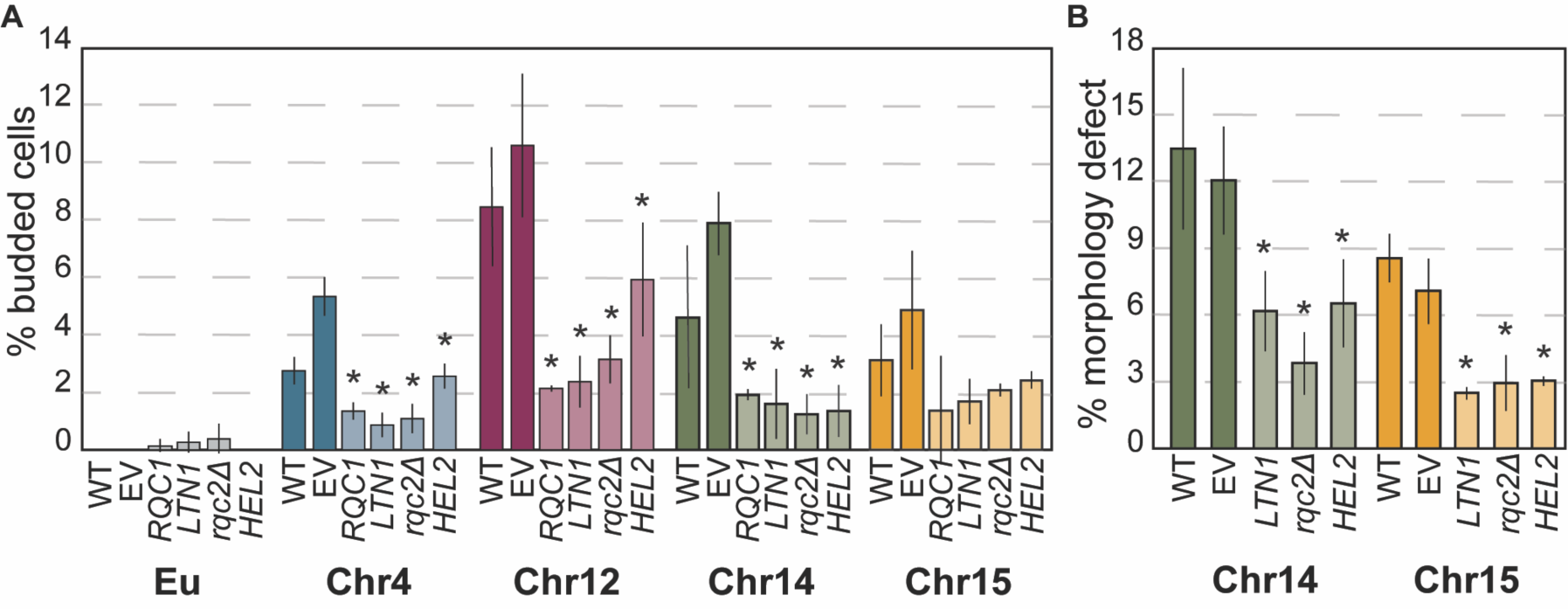
Genetic perturbation of the RQC pathway modulates cell-cycle defects. (A) Percent budded cells in euploid (Eu) and aneuploid cultures harboring empty vector (EV) or plasmids encoding *RQC1*, *LTN1*, or *HEL2* or in a strain lacking *RQC2* (*rqc2τι*) after 2 days of culturing. (B) Percent cells with morphology defects YPS1009_Chr14 and _Chr15 after 2 days. Asterisk, p < 0.05 comparing strains with gene plasmids to EV or *rqc2τι* to WT; n = 3, paired T-test.

### Aneuploid cells accumulate stalled ribosomes and RQC defects

Results above point to a problem in RQC during quiescence in aneuploids. We therefore quantified RQC defects using a reporter with a stall-inducing stretch of 12 arginine codons between GFP and tdTomato coding sequences (**Fig 3A**). Healthy cells can readily dismantle and degrade ribosomes stalled on the reporter; we confirmed that euploid cells lacking Rqc1 or Ltn1 accumulate a smeared product consistent with CATylated GFP, as well as full-length readthrough protein (**Fig 3B**). As expected, the absence of Rqc2 in euploids produced a crisp GFP band consistent with the absence of CATylation on the stalled product; surprisingly the *rqc2Δ* mutant displayed very little full-length product, implicating a role for Rqc2 in translational readthrough.

**Figure 3.**
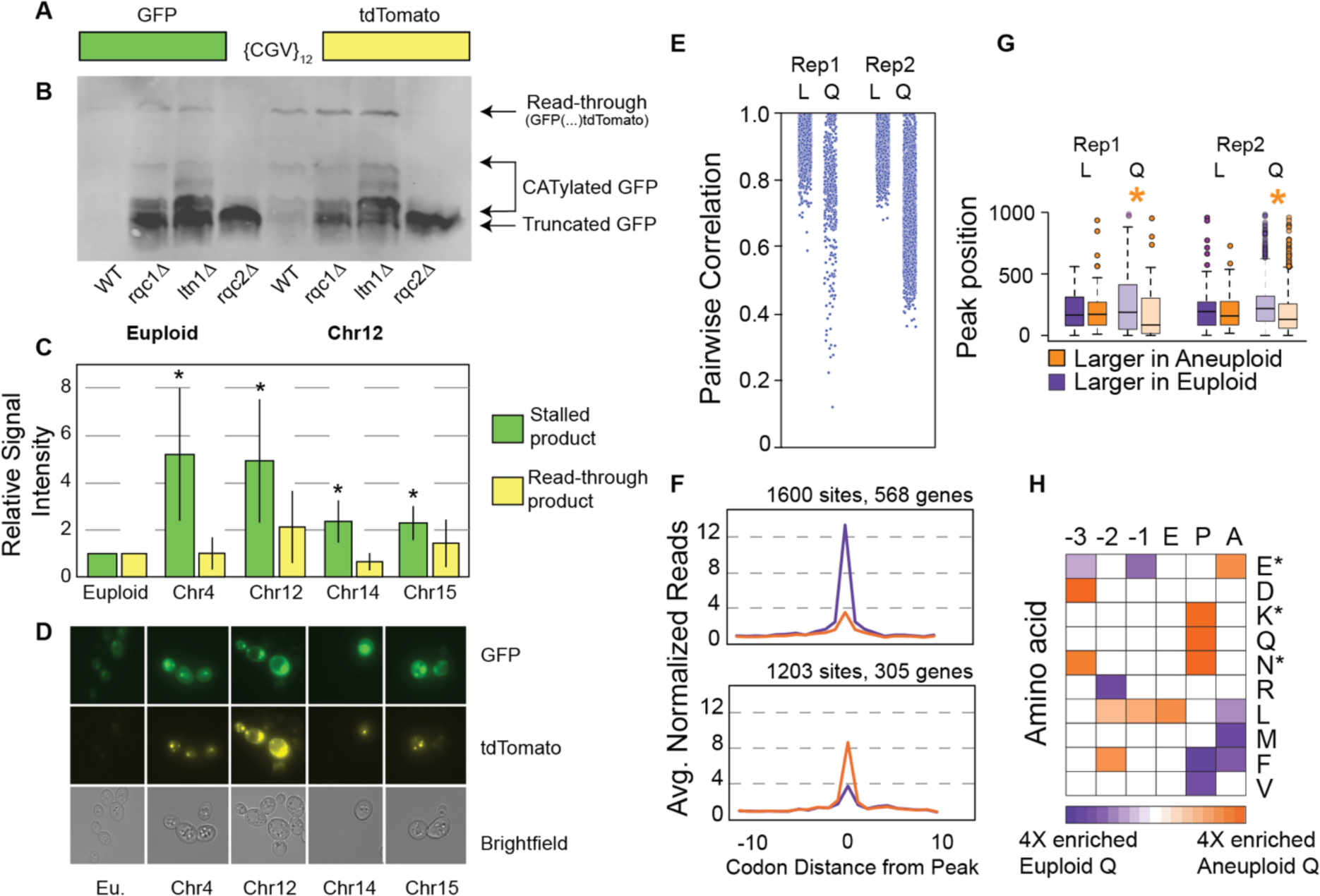
Aneuploids show signs of RQC defects. A) Ribosome stalling reporter, see text. B) Anti-GFP Western blot of euploid and YPS1009_Chr12 RQC mutants carrying the ribosome stalling reporter, cultured for 4 days. C) Relative signal intensity of normalized RQC defect products versus euploid. Asterisk, p<0.05, T-test. D) Representative microscopy images showing GFP (top), tdTomato (middle), or bright field images. E) Pairwise correlations in traces of normalized ribosome occupancy for transcripts measured in euploid and YPS1009_Chr12 aneuploid cells in log (L) or quiescent (Q, day 4) phase. F) Average ribosome occupancy (read count at each codon normalized to gene-body counts, see Methods) for peaks significantly with higher read count in euploids (purple, top) or in YPS1009_Chr12 (orange, bottom, FDR < 0.05). G) Distribution of codon positions for peaks with higher read count in euploids (purple) or in aneuploids (orange). Asterisk, p<0.05, Wilcoxon test. Some outliers are omitted from display. H) Enrichment of amino acids (rows) encoded at each position (columns) relative to the ribosome P site. Asterisk, enrichments seen previously in yeast ribosome stall sites^44^. Enrichments are only shown for positions with statistically significant differences in amino acid frequency (FDR < 0.05, Fisher’s exact test).

In contrast to euploid cells, all of the wild-type aneuploids tested accumulated significantly more CATylated GFP as well as read-through product after 4 days of culturing (**Fig 3B-C**). Furthermore, all of the aneuploids showed bright fluorescent signal by microscopy, along with foci containing GFP and tdTomato that implicate aggregation (**Fig 3D** and below). Interestingly, in many of the aneuploid cells the full-length product localized closely to the nuclear signal, especially YPS1009_Chr12. This was intriguing, since defective translation products can accumulate in the nucleus when not degraded^85–87^.

To investigate ribosome stalling on native transcripts, we performed replicate ribosome footprinting on euploid and the YPS1009_Chr12 aneuploid, in log phase and after 4 days of culturing. Globally, ribosome profiles from exponentially growing cells were remarkably similar: most traces of normalized ribosome occupancy were highly correlated comparing individual transcripts in euploid versus aneuploid cells in log phase (**Fig 3E**). In contrast, ribosome occupancy traces for many transcripts were poorly correlated in quiescent samples in both replicates. To further investigate, we identified ribosome pause sites (occupancy ‘peaks’) that were more or less prominent in quiescent aneuploid cells versus euploid (FDR < 0.05, Fisher’s exact test, see Methods). We identified 1600 sites in 568 transcripts that displayed higher ribosome occupancy in quiescent euploid samples and 1203 sites in 305 transcripts with higher occupancy in the quiescent aneuploid (**Fig 3F**). Interestingly, many transcripts showed multiple peaks on the same mRNAs with opposing effects, where some occupancy peaks were higher in aneuploids but other peaks were larger in euploids. While investigating this, we noticed that many aneuploid-enhanced peaks were nearer to the start of the transcript (**supplemental Fig 3**); in fact, the set of peaks more prominent in the aneuploid occurred closer to the start codon than the set of peaks with higher occupancy in the euploid (**Fig 3G**, p<0.05, Wilcoxon rank-sum test). While future investigation will be required, this signature may reflect feedback from stall sites to slow upstream translation initiation or elongation^88–90^.

Previous studies show that ribosome stalling occurs at specific residues^44,91–94^. We wondered if peaks in ribosome occupancy that differed in aneuploid versus euploid cells differed in sequence characteristics. We compared the frequency of encoded amino acids surrounding differential peaks. Aneuploid-enhanced peaks were enriched for specific signatures previously identified at age-induced sites of ribosome stalling in yeast (**Fig 3H**), including lysine or asparagine at the ribosome P-site and glutamate at the A-site^44^. Interestingly, aneuploid-enhanced peaks were also enriched for leucine codons at several sites encoded before the peak. Importantly, peaks that were called significantly different between log-phase aneuploid compared to euploid cells were typically subtle in magnitude (**supplemental Fig S3B**) and showed no significant difference in associated sequences (FDR > 0.05 at all sites). Thus, aneuploid YPS1009_Chr12 cells have substantially different ribosome occupancy peaks – but only during quiescence phase – consistent with differences in ribosome stalling.

#### Inducing ribosome stalling in euploids disrupts quiescence and lifespan

If problems managing ribosome stalling drive quiescence defects in aneuploids, we hypothesized that increasing the level of ribosome stalling in euploids would cause similar defects. Indeed, this was the case. Low doses of NTC induce ribosome stalling, evident by increased RQC intermediates from the stalling reporter (**Fig 4A**). This dose of NTC was enough to significantly increase the number of euploid cells that remained budding at day 3 (**Fig 4B**). Remarkably, simply over-expressing the RQC reporter had the same effect, while expressing the reporter in conjunction with NTC treatment significantly exacerbated the defects, much beyond a sodium chloride-stress control (**Fig 4B**). Increasing the levels of ribosome stalling in the euploid also slightly but significantly disrupted cell densification compared to the euploid control (**supplemental Fig S4**). In contrast, deleting *RQC1* or *LTN1* without increasing stalling produced only a mild arrest defect, demonstrating that increased stalling is important for the phenotype. Finally, activating ribosome stalling in euploid cells significantly shortened lifespan to much greater levels than a salt-stress control (**Fig 4C**). Together, these results show that it is the increase or perhaps quality of ribosome stalling that affects quiescence and lifespan, rather than a defect managing basal levels of stalling seen in the euploid (see Discussion).

**Figure 4.**
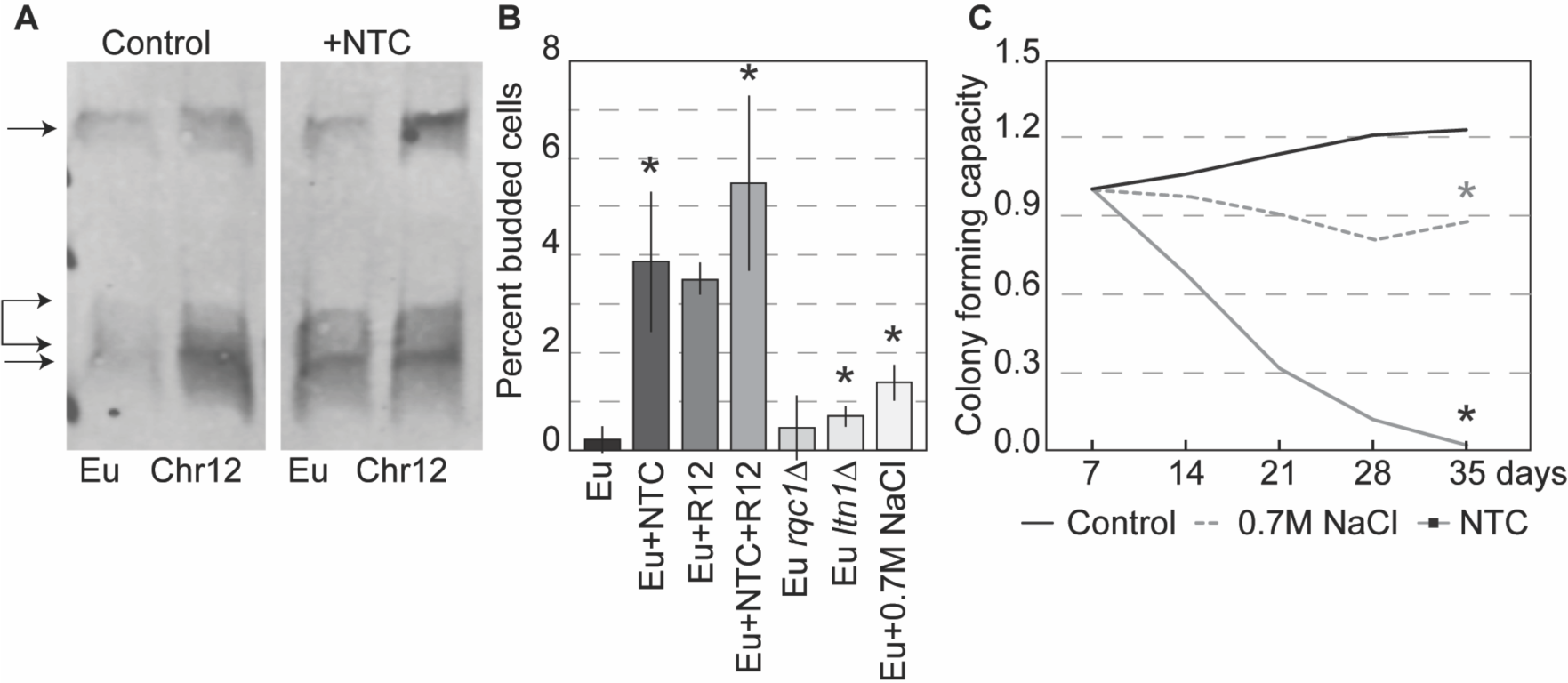
Induction of ribosome stalling disrupts quiescence and lifespan in euploid cells. (A) Representative anti-GFP Western blot of euploid (Eu) and YPS1009_Chr12 cells with the stalling reporter, as annotated in Fig 3B, cultured for 4 days with and without NTC. (B) Percent budded cells in euploid cells treated with 0.7M NaCl, exposed to low-dose (1μg/mL) NTC, carrying the ribosome stalling reporter (“R12”), or both NTC treatment and R12 reporter at 3 days. Asterisk, p < 0.05 compared to untreated control (n = 3), paired T-test. (C) Average colony forming capacity of YPS1009 euploid cells untreated (control) or treated with low-dose (1μg/mL) NTC or 0.7 M NaCl. Asterisk, p<0.05 compared to untreated control at 35 days, n = 4; T-test.

Interestingly, we noticed that NTC treatment and/or the stalling reporter significantly exacerbated defects seen in aneuploid cells, including odd morphologies of starved cells entering quiescence. In fact, NTC treatment of YPS1009_Chr12 induced a small number of bi-lobed cells characteristic of YPS1009_Chr14 entering quiescence, and in a few instances produced cells in which nuclear division occurred perpendicular to the division plane (**supplemental Fig S5A**). These morphologies are reminiscent of those caused by defective Cdc34, the E2 ubiquitin conjugase of the SCF complex that marks cell-cycle regulators for timed degradation by the ubiquitin-proteasome system (UPS)^95–97^.

#### Genes linked to ubiquitin metabolism alter aneuploid arrest phenotypes

To further dissect how RQC defects could impact cell cycle arrest, we returned to hits from our screen in YPS1009_Chr12 as a representative aneuploid. Several genes from the screen were linked to SCF-dependent protein degradation, including Cdc34 regulator *UBS1*^98,99^, chaperone *SGT1* that associates with SCF^100^, and *POG1* that has been implicated in G1/S regulation and can suppress defects in E3 ubiquitin ligase Rsp5^101–103^. Duplication of all three genes partly alleviated the arrest defect in YPS1009_Chr4 and/or YPS1009_Chr12 **(Fig 5)**; we also tested SCF-associated F-box subunit *GRR1,* which produced a mild improvement but missed the significance threshold.

**Figure 5.**
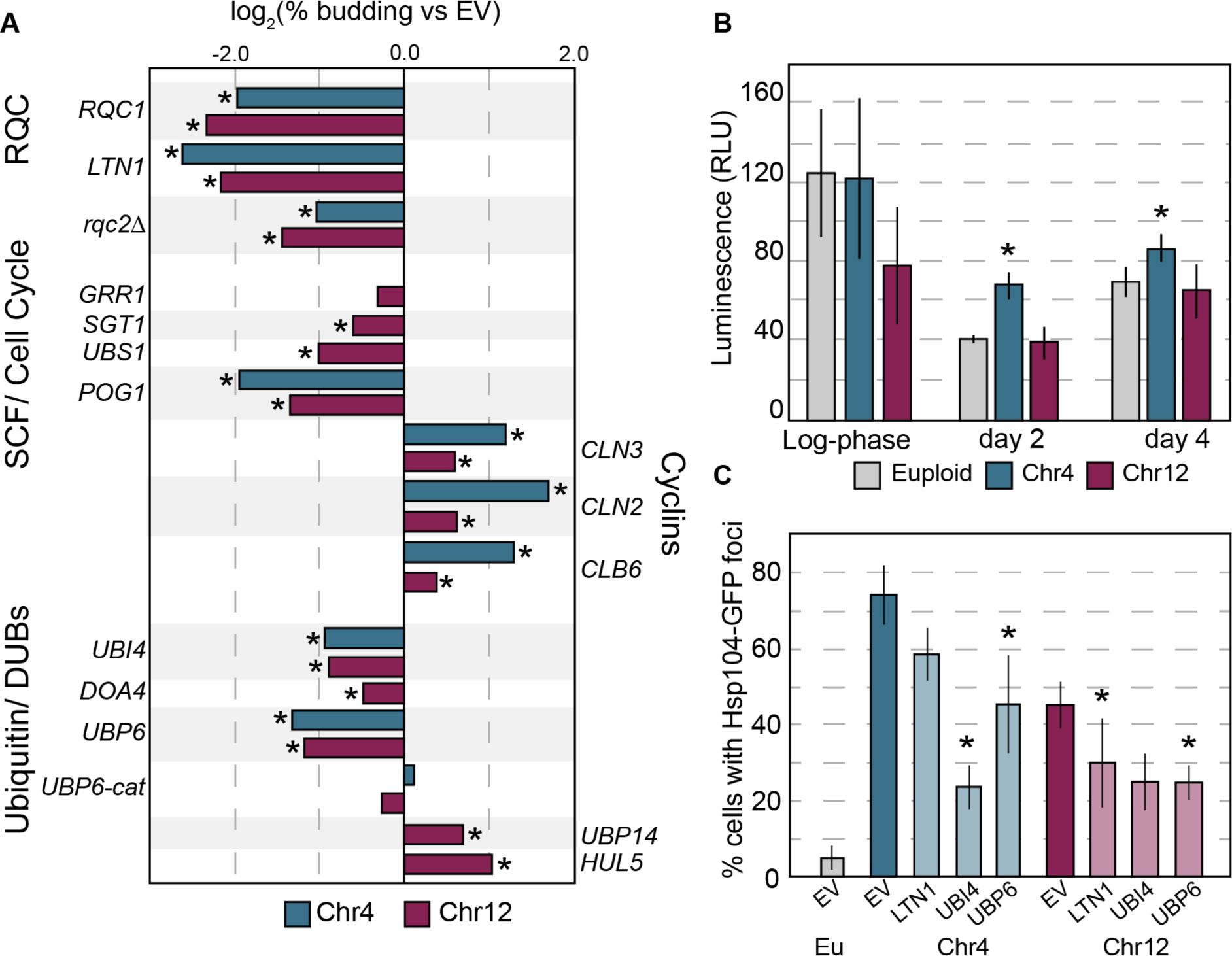
Genetic perturbation influences aneuploidy phenotypes. A) log_2_ change in % budded cells for Chr4 (blue) or Chr12 (magenta) aneuploids harboring plasmids expressing different genes. Other tested genes had no significant effect (*AFG3, ATG12, OTU1, PCL2, PRE1, RPN11, YGP1, LEE1)*. Not all genes were tested in YPS1009_Chr4. B) *In vitro* chymotrypsin-like proteasome activity, determined by cleavage of a luminescent substrate, in euploid and aneuploid lysates. Asterisk, p < 0.05 compared to euploid control, n=3; paired T-test. C) Percent cells with Hsp104-GFP foci (n=3) for Chr4 or Chr12 aneuploids harboring indicated gene duplications. Asterisk, p<0.05 compared to empty vector control.

We hypothesized that a defect in SCF may lead to aberrant cyclin abundance. In fact, YPS1009_Chr12 grown for 1 or 2 days showed visible accumulation of HA-tagged Cln3 products, whereas euploid cells did not (**supplemental Fig S5D**). Cln3 catalyzes cell-cycle entry, in part by triggering nuclear eviction of transcriptional repressor Whi5 to enable induction of S-phase genes^104^. We found that 95% of budded YPS1009_Chr12 cells lacked nuclear Whi5-GFP, consistent with inappropriate entry into the cell cycle despite undetectable glucose in the culture (**supplemental Fig S5B-C**). We reasoned that, if cyclin degradation is defective in aneuploids then over-expression of cyclins may exacerbate phenotypes. Indeed, we found that duplicating *CLN3* or downstream G1 and S phase cyclins *CLN2* or *CLB6* substantially increased budding in both YPS1009_Chr4 and YPS1009_Chr12 aneuploids, with only a weak effect on euploid cells (**Fig 5A** and **Fig S5E**). These results strongly suggest that cyclins including Cln3 are not properly degraded in aged aneuploids and show that increased cyclin gene copy exacerbates defects.

One possibility is that chromosome amplification disrupts proteasome function, as previously proposed in the aneuploidy-sensitized laboratory strain^105,106^. However, this was not the case in the wild strain background: we measured proteasome activity in cell lysates using luminescent reporters that do not require ubiquitination for degradation^107^. Aged aneuploids showed similar proteasome activity to euploid cells, for all three protease activities tested (**Fig 5B** and **supplemental Figure S6**, see Methods). Thus, defects in proteasome activity *per se* do not explain quiescence defects in aneuploid cells.

However, we found that duplication of the stress-induced polyubiquitin gene *UBI4* significantly alleviated the aneuploid arrest defect in both aneuploids tested (**Fig 5A**). Likewise, duplication of two of the major deubiquitinases (DUBs) important for ubiquitin recycling, *UBP6* and *DOA4*, improved cell-cycle arrest (**Fig 5A**). This is the opposite effect reported for the sensitized W303 strain, where deletion of *UBP6* provided a benefit, reportedly by relieving proteasome inhibition that occurs through a separable Ubp6 domain^105,106,108^. To distinguish between these functions, we tested catalytically inactive ubp6-C118A that can still inhibit proteasomal processivity but lacks ubiquitin recycling activity. This mutant did not mitigate aneuploid arrest defects, indicating that the deubiquitinase activity is required (**Fig 5A**). In contrast, duplicating ubiquitin ligase *HUL5*, which antagonizes ubiquitin recycling by extending ubiquitin chains and increasing proteasome processivity, exacerbated arrest defects (**Fig 5A**). Duplication of a different deubiquitinase *UBP14,* which disassembles unanchored ubiquitin chains and specific targets^109^, increased defects. These results suggest that it is aneuploidy-specific dysfunction in ubiquitin metabolism and not proteasome activity *per se* that underlies aneuploid defects. Notably, none of the gene duplications tested alleviated the arrest defect to the same level as *RQC1* or *LTN1*.

#### Proteostasis stress is alleviated by duplication of RQC or ubiquitin

Trisomy 21 in humans and chromosome amplification in sensitized W303 yeast is associated with increased protein aggregation, although the source of proteostasis stress is not known. We previously showed that wild aneuploid yeast strains do not show signs of proteostasis dysfunction unless stressed by *SSD1* deletion or treated with translational inhibitor NTC^13,22^. Here we found that aging also induces protein aggregation in wild aneuploids: at 7 days of culturing, 50-70% of aneuploids with extra Chr4 or Chr12 harbored foci of protein disaggregase Hsp104, compared to ∼5% of euploid cells. The proportion of aneuploids with such aggregates decreased substantially upon duplication of RQC gene *LTN1*, polyubiquitin *UBI4*, or deubiquitinase *UBP6* (**Fig 5C**). *UBI4* provided a greater benefit to YPS1009_Chr4 compared to YPS1009_Chr12, while *LTN1* produced a bigger effect in YPS1009_Chr12 cells. Interestingly, *UBI4* is encoded on Chr12 and thus already duplicated in this aneuploid strain, whereas *RQC1* is encoded on Chr4, raising the possibility of some natural protection for these aneuploids. Nonetheless, the fact that *LTN1* over-expression reduces protein aggregation strongly suggests that defects in the RQC pathway are an unrecognized source of proteostasis stress in aneuploid cells (see Discussion).

We considered several models for how RQC defects could perturb ubiquitin homeostasis. One possibility is that protein aggregation triggered by RQC defects depletes free ubiquitin in aged aneuploids, especially since non-mitotic cells rely on ubiquitin recycling for proteostasis maintenance^110^. Yet, Western blots of bulk-culture lysates revealed that monoubiquitin is still visible compared to the euploid. We next asked if ubiquitin localization was different in aneuploid cells using fluorescence microscopy. Indeed, the ubiquitin profile was markedly different in aged YPS1009_Chr4 and _Chr12 aneuploids (**Fig 6A**). Whereas ubiquitin was evenly distributed across most of the aged euploid cells, the majority of cells with extra Chr4 or Chr12 showed very bright ubiquitin signal, often in discrete puncta (**Fig 6A-B**). YPS1009_Chr4 had especially high ubiquitin levels in these puncta. Together, these results are consistent with aggregates of ubiquitinated proteins (see Discussion).

**Figure 6.**
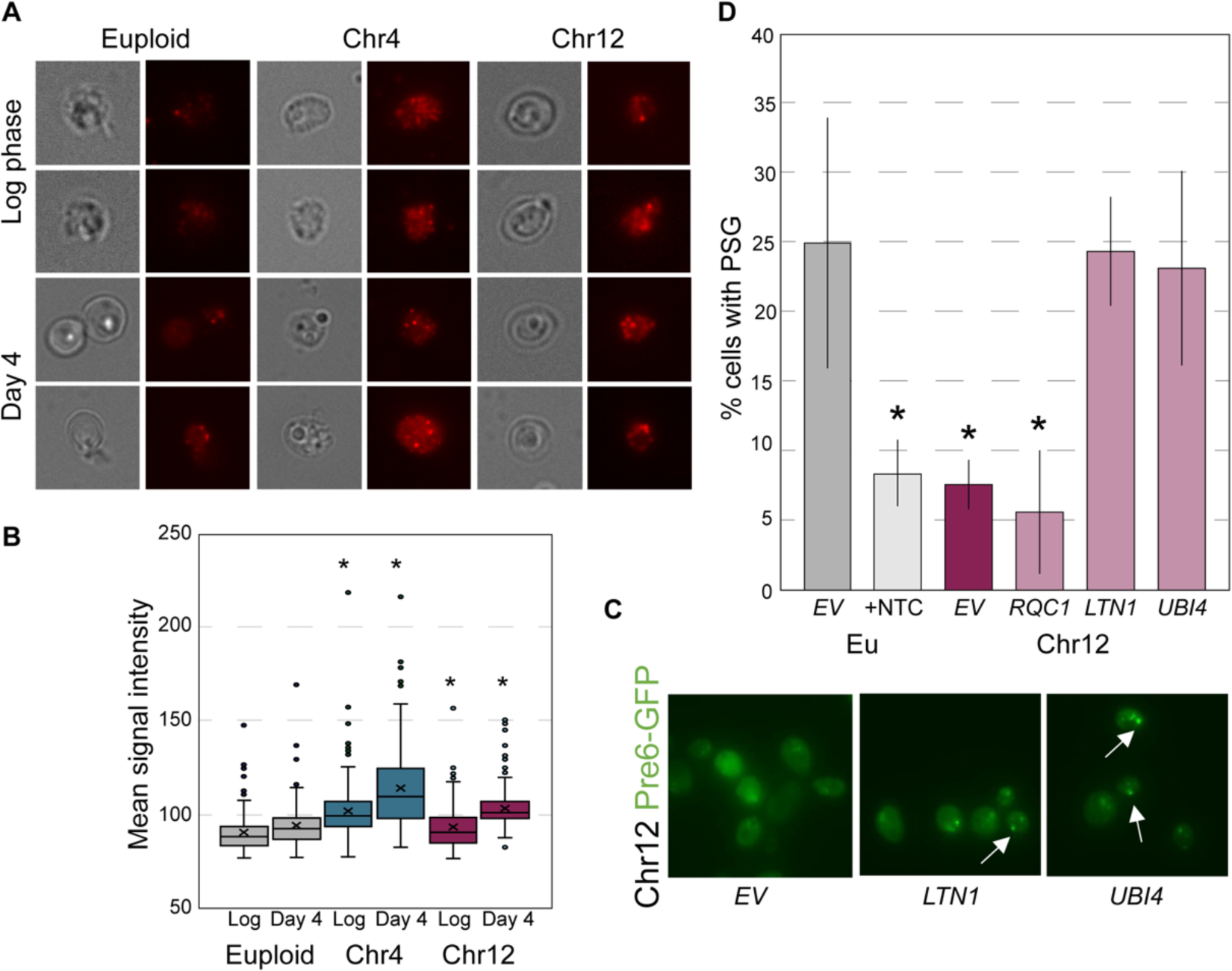
Aneuploids exhibit altered ubiquitin distributions. A) Representative brightfield and fluorescent images of euploid and aneuploid cells at log-phase and day 4 stained with fluorescent anti-ubiquitin antibody. B) Mean intensity of ubiquitin signal per cell in euploid and aneuploid cells at log-phase and day 4. Asterisk, p<0.05, Wilcoxon rank sum test. C) Representative images of Pre6-GFP foci in YPS1009_Chr12 carrying indicated plasmids. Arrows highlight PSGs. D) Percent of cells with Pre6-GFP foci, in euploid (Eu) and YPS1009_Chr12. Asterisk, p<0.05, n=3, T-test compared to euploid control.

We also considered how altered ubiquitin distribution might influence proteasome localization. Upon quiescence entry, proteasomes move from the nucleus to cytosolic foci called Proteasome Storage Granules (PSGs). PSGs are thought to form around free ubiquitin in the cytosol, and they are important for longevity and quiescence exit^111,112^. We found that cells carrying an extra Chr12 had a significant defect forming PSGs, since few cells formed foci of proteasome subunit Pre6-GFP (**Fig 6C-D**). Once again, over-expression of *LTN1* or *UBI4* recovered the defect in YPS1009_Chr12 cells; although *RQC1* complemented the budding defect, it did not correct PSG formation for reasons that are not clear. The YPS1009_Chr4 aneuploid did not show a defect compared to the euploid, although increasing *LTN1* or *UBI4* increased PSG formation in many cells (**supplemental Fig S7**). Importantly, treating the euploid cells with NTC that induces ribosome stalling blocked PSG formation in most cells (**Fig 6D**). These results are consistent with the model that RQC dysfunction disrupts ubiquitin stasis, produces protein aggregates, and influences PSG formation in multiple strains (see Discussion).

## DISCUSSION

Premature aging is a hallmark of Down syndrome (DS) and has also been observed in aneuploidy-sensitized laboratory yeast^2,61^. Here we delineate that multiple features of premature aging, including defects in quiescence, mounting proteostasis stress, and shortened lifespan, result from a generalizable consequence of chromosome amplification in yeast, across genetic backgrounds and amplified chromosomes. Together with shared signatures of other aneuploidy syndromes, including premature senescence and protein aggregation in human trisomy 13 and 18^113,114^, this strongly implicates premature aging as a conserved consequence of aneuploidy across species. Despite prior indications of accelerated aging in people with DS, the mechanistic basis has been a mystery. Our results in yeast show that defects in translation and ribosome quality control contribute: aneuploid yeast strains accumulate RQC intermediates, show aberrant ribosome profiles, and harbor aneuploidy-associated protein aggregates. Several of these aneuploidy phenotypes, including defective cell-cycle arrest and protein aggregation among others, can be partly corrected simply by increasing abundance of RQC subunits. In contrast, inducing ribosome stalling in euploid yeast produces similar aberrations in aging and lifespan, confirming a causal link. These results are interesting given that healthy aging is already associated with a decline in both translational fidelity and proteostasis management^45,115,116^. Our work here adds to a growing body of evidence that chromosome amplification accelerates that decline.

Integrating our results and others presents a model for how aneuploidy catalyzes these effects (**Fig 7**). Most transcripts encoded on the amplified chromosome are expressed at higher abundance than in euploids, although some genes show muted expression compared to DNA content^14,15,117,118^. However, many of the encoded proteins are not elevated to the same extent in yeast or human cells^106,119–123^. In both yeast and trisomic human fibroblasts, this is due to increased turnover of proteins encoded by duplicated genes^120,122,124^, which minimizes their abundance differences even when underlying mRNA levels are elevated. This result strongly suggests that over-abundant mRNAs are still translated in aneuploid cells. We propose that the increased translational burden caused by an over-abundance of translated mRNAs, coupled with a natural decline of translational fidelity with age, depletes limiting translational quality control proteins. Since several proteins in the RQC pathway are stoichiometrically limiting^36,80,81^, increasing the molecular mass of stalled ribosomes would quickly overwhelm the pathway. The consequences may be two-fold: on the one hand, producing an accumulation of RQC intermediates when the pathway is activated, and on the other hand, causing mistranslation and frameshifting when the pathway is not activated. Both scenarios catalyze protein aggregation, which can further accumulate due to age-associated proteostasis decline. This hypothesis is consistent with our results, since over-expressing RQC activator Hel2 or downstream RQC effectors Rqc1 and Ltn1 are both beneficial in aneuploids. That we were unable to make or maintain aneuploid cells that lack *HEL2* underscores the importance of the RQC pathway – but even when the pathway is activated, RQC intermediates accumulate.

**Figure 7.**
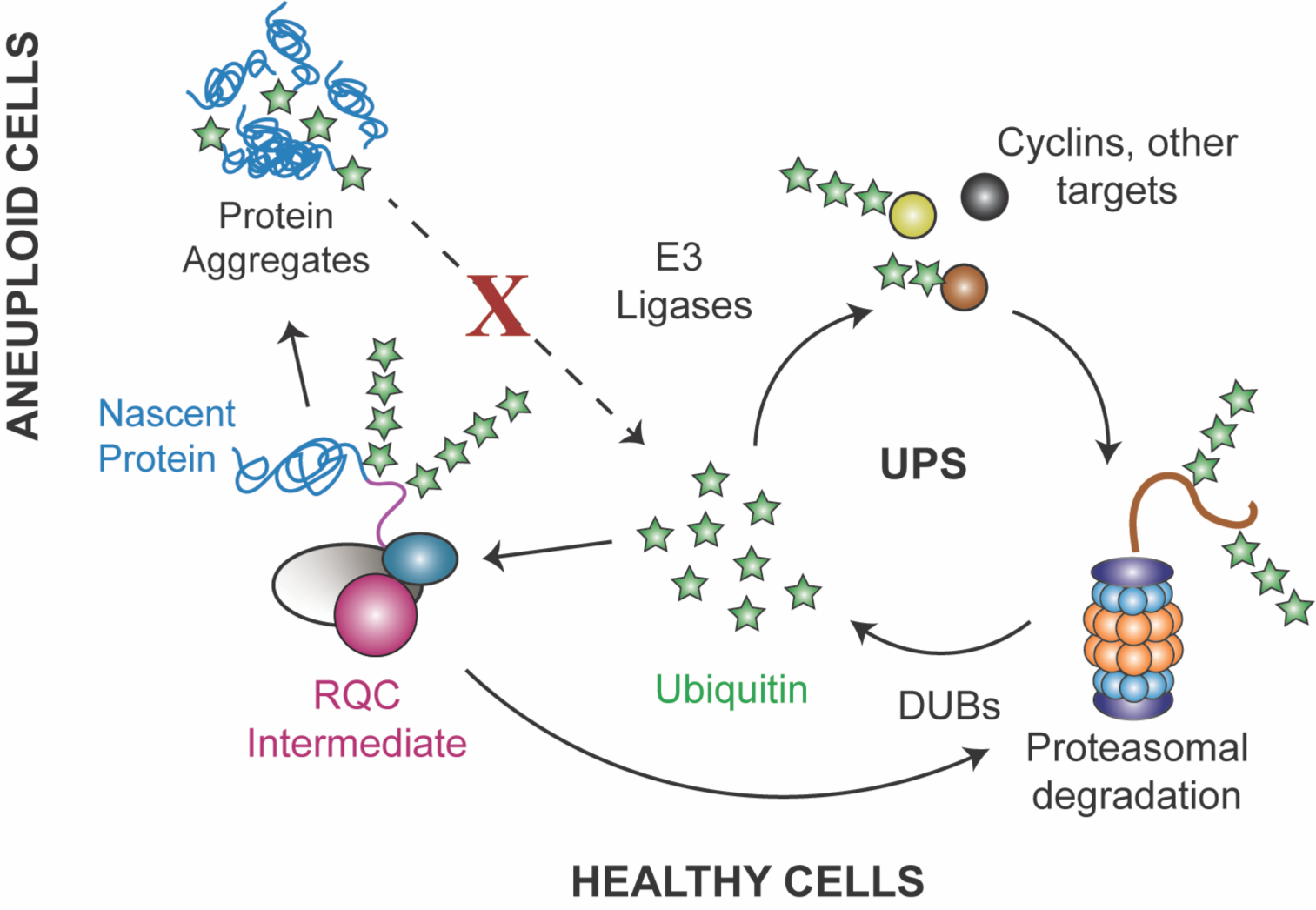
Model for RQC impacts on aging in aneuploid cells. Healthy cells maintain a balance or ubiquitin stasis as part of the Ubiquitin Proteasome System (UPS). We propose that defects in RQC produce protein aggregates that sequester ubiquitin, thereby depleting the pool of accessible ubiquitin, see text for details.

Our proposed model is that defects in quiescence and aging manifest when RQC dysfunction promotes protein aggregation that sequesters ubiquitin from critical functions. Somewhat surprisingly, aneuploid cells studied here have no defect in proteasome activity – rather, we argue that proteostasis stress emerges from a defect in ubiquitin stasis that alters the landscape of ubiquitinated proteins. Several aging signatures, including defective arrest and accumulation of Hsp104 foci, were alleviated simply by increasing ubiquitin levels or ubiquitin recycling. Dividing cells can likely manage the stress of aneuploidy, relying on growth-coupled ubiquitin synthesis along with mitotic division to retain damaged proteins in the mother cells so that daughter cells are born anew^125–128^. But post-mitotic quiescent cells must rely on alternate mechanisms, including ubiquitin recycling and aggregate turnover^110^. In fact, quiescent yeast cells are heavily dependent on DUBs to maintain ubiquitin levels and ubiquitin-dependent processes^125^. A shortage of accessible ubiquitin could significantly alter the ubiquitination landscape. It is worth noting that ubiquitin metabolism is especially important for translational fidelity: yeast mutants defective in ubiquitin metabolism are sensitive to translational inhibitors including those in the same class as NTC^129–131^. Thus, inaccessibility of ubiquitin could further exacerbate translational defects in aneuploid cells.

Although free ubiquitin appears not to be completely depleted in analysis of bulk cultures, it may be limiting in subsets of cells. In fact, many of the phenotypes we investigated show cell-to-cell heterogeneity, *e.g.* in the proportion of cells that fail to arrest (**Fig 1**), the number of cells with substantial Hsp104 foci (**Fig 5**), or the variation in quantity and quality of ubiquitin distributions (**Fig 6**). This observation suggests that cells may operate around a threshold of biosynthesis errors, below which they can function but over which they experience cellular collapse^13^.

Aneuploid cells may be closer to that threshold, such that stochastic fluctuations push many individual cells past the point of return to a healthy state. This could contribute to previously observed heterogeneity in aneuploid phenotypes ^132,133^. One of the signatures of the cellular collapse may include protein aggregation and ubiquitin sequestration in aggregates. In fact, ubiquitin sequestration is prominent in neurodegeneration and diseases including Huntington’s, Alzheimer’s, and ALS^134–137^. Notably, these diseases have all been linked to RQC defects^50,138–140^.

An altered ubiquitin landscape could explain other peculiarities, including chromosome-specific effects. Despite a generalizable contribution of aneuploidy to premature aging, there are obvious chromosome-specific phenotypes, including quiescence-induced morphology defects for Chr14 and Chr15 amplification, severe densification abnormalities for Chr4 and Chr12 duplication, and chromosome-specific differences in lifespan. These defects are likely influenced by specific genes on each chromosome, but notably they are all revealed when cells transit to quiescence. One explanation may be the interplay between chromosome-specific gene amplification and competition among different E3 ligases for limiting ubiquitin. The *S. cerevisiae* genome encodes 24 DUBs and at least 85 ubiquitin E3 ligases, whereas humans encode >100 DUBs and >600 E3s^141,142^. Many of these enzymes target specific proteins and ubiquitin moieties, across different cell types and subcellular regions, giving individual DUBs and E3 ligases specificity in the processes they influence^143,144^. Strain-specific differences in proteome content may influence competition for ubiquitin and ubiquitinated proteins by E3 ligases and DUBs, thereby producing chromosome-specific phenotypes from a generalizable defect in ubiquitin availability. It is notable that a variety of DUBs and E3 ligases have been implicated in aneuploidy, in sensitized W303 yeast^24,61,105,106^ and DS, including such genes amplified on human chromosome 21^33,145,146^.

Our results indicate that defects in RQC contribute to hallmarks of premature aging. Increasing the abundance of RQC subunits reduced several aging signatures; however, it did not prolong aneuploid lifespan. This is consistent with the results of our screen, where RQC and ubiquitin regulators produced a benefit early in the aging time course, whereas a different set of genes, including many linked to mitochondrial functions, were enriched later. Mitochondrial function is known to be important for yeast lifespan, and it is defective in both yeast and DS aneuploids^13,147^. It is also possible that aged aneuploid yeast suffer from autophagy defects, as previously observed in sensitized W303 and DS cells^11,148,149^, a defect that could also explain why ubiquitin aggregates are not turned over (**Fig 6C**). Nonetheless, improving RQC and ubiquitin availability clearly impacts healthspan of aneuploids, and could contribute to lifespan extension in the context of other improvements in metabolism or autophagy. An exciting avenue for future work is to test if the conserved impact of aneuploidy on aging is due to conserved mechanisms related to RQC.

## METHODS

### Strains and Plasmids

Strains used in this study are listed in Table 1. YPS1009 aneuploids were generated in Rojas et al.^23^ Most chromosome duplications are stable for many generations; maintenance of aneuploidy was confirmed periodically by plating cultures from each experiment onto synthetic complete media with selection for marked chromosomes (SC -HIS +NTC). Gene deletions were generated by homologous recombination of the Hph-MX drug resistance cassette into the designated locus, followed by diagnostic PCR to confirm correct integration and absence of the target gene. Aneuploidy was confirmed and periodically checked through diagnostic qPCR of one or two genes on the affected chromosomes normalized to a single-copy gene elsewhere in the genome – normalized ratios close to 2 reflect gene duplication, and ratios between 1.2–1.8X indicated partial loss of aneuploidy in the cell population. *HSP104*-GFP was generated by integrating a GFP-ADH2 terminator::HIS3 cassette into the *HSP104* locus^150^ via homologous recombination. Aneuploid Hsp104-GFP strains were then generated through mating and dissection, crossing AGY1970 to relevant aneuploids. The GFP-{CGV}12-tdTomato stalling reporter was generated by PCR sewing and cloning the generated fragment into a KAN-marked CEN plasmid (pJH2). Unless otherwise noted, plasmids used in this work were from the MoBy 1.0 plasmid library^64^.

### Growth conditions

Unless otherwise noted, all experiments were performed in rich YPD (Yeast extract, Peptone, Dextrose) medium. Quiescence cultures were generated by inoculating liquid YPD medium at an optical density (OD_600_) of 0.05. Cultures were allowed to reach saturation and then maintained at 30°C in a shaking incubator for the number of days indicated, with no nutrient supplementation. Maintenance of aneuploidy was verified by plating an aliquot of aneuploid cultures onto rich YPD plates, then replica plating to SC -HIS +NTC after 24 hours to determine maintenance of the two chromosome markers. Aneuploidy was also periodically verified through diagnostic qPCR as described above. Where indicated, cells without the NAT-MX resistance cassette were treated with 1 ug/mL NTC after cells reached mid-log phase.

### Microscopy

#### Bud indexing

Cultures were grown in YPD for 2 days and fixed with formaldehyde as described previously^151^. Cells were stained with DAPI using NucBlue ReadyProbes (ThermoFisher, R37606) for 20 min at room temperature to stain DNA. Images were acquired as z-stacks every 0.2 mm using an EVOS FL Auto 2 with a 100x Nikon oil immersion objective equipped with an EVOS DAPI light cube. Cells were scored as budded or unbudded based on morphology and DAPI signal. Budded cells were scored as cells with budded morphology, lack of septum, and a bud either lacked DAPI signal (S-phase to early G2 buds) or possessed bar nuclei (late G2 to M-phase buds). Late-stage buds were likely undercounted by this method. A minimum of 5 diverse xy positions and 100 cells were scored per replicate. A minimum of 3 biological replicates were conducted. Bud index was calculated as a percent cells scored as budded ((# of budded cells/ # of cells scored) * 100). Indexing was conducted in the same way for indicated strains carrying plasmids, except cells were grown in YPD with G418 to select the plasmids.

#### Live cell microscopy

After culturing for 4 days, live cells were deposited onto plain glass slides. Images were acquired as z-stacks every 0.2 mm using an EVOS FL Auto 2 with a 100x Nikon oil immersion objective. GFP and tdTomato images were acquired with EVOS GFP and RFP light cubes, respectively. Fluorescent images represent collapsed Z-stacks, and brightfield images represent one z-plane.

#### Immunofluorescence

Cells were harvested during exponential phase (OD_600_ 0.4 - 0.6) or 4 days after start of culturing and fixed with 4% formaldehyde for 15 min at 30C followed by centrifugation. Cells were spheroplasted with zymolyase then treated with 0.1% SDS in buffer A (100 mM Tris, pH 8 1M sorbitol) for 10 min. After washing with buffer A, cells were then plated onto a 96-well black-walled plate with a poly-L-lysine coated coverglass bottom (Cellvis). After 30 min of incubation with blocking buffer (50 mM Tris pH 8, 150 mM NaCl, 1% nonfat dry milk, 0.5 mg/ml BSA, 0.1% Tween 20), cells were exposed to anti-ubiquitin antibody (Milipore Sigma, MAB1510) in blocking buffer overnight at 4C. After washing with blocking buffer, cells were exposed to anti-mouse Alexa Fluor 647 antibody (Life Technologies, A21235) for 1 hour at room temperature. After washing with blocking buffer, ProLong Gold Antifade Mountant (ThermoFisher, P36934) was applied to each well. Images were acquired as Z-stacks every 0.2 mm using an EVOS FL Auto 2 with a 100x Nikon oil immersion objective. FIJI (imageJ) was used to determine ubiquitin signal intensity. Brightfield images were used to generate cell masks and mean signal intensity of ubiquitin immunofluorescence was computed for each cell.

### Cell-density fractionation

Density gradient sedimentation and fractionation of stationary phase cultures was performed using Percoll (Sigma, P1644) as described previously^152^. Gradients were split evenly between two fractions. Fractions were collected using 18-gauge needle and 10 ml syringe, harvesting the heavy fraction first, then washed once in PBS, and resuspended in 1 mL of PBS. Fractions were quantified using a hemacytometer.

### Chronological lifespan assay

Chronological lifespan was determined by plating for colony forming capacity over time. At various times over long-term growth, an aliquot of culture was harvested, OD_600_ measured, and cells diluted serially to a 40,000X dilution, which was spread on YPD plates. Plates were incubated for 48 hours, and viable colonies were counted using ImageJ Colony Counter Plug-in (ImageJ) to quantify colony forming units. Colony forming capacity was calculated as colony forming units divided by optical density measured at the indicated day.

### RNA sequencing

RNA-seq was performed using total RNA isolated from log-phase and quiescent cultures. Cultures were started at OD_600_ 0.05. Log-phase cultures were harvested after precisely three generations. Day 1, 3 and 7 cultures were harvested 24, 72, and 168 hours after log-phase cultures were harvested. OD-normalized samples were pelleted by centrifugation and flash frozen with liquid nitrogen and maintained at −80°C until RNA extraction. Samples were mixed with a defined number of flash-frozen *Sz. pombe* cells before RNA extraction, to later serve as per-cell normalization. Total RNA was extracted by hot phenol lysis^153^. Mechanical disruption was required to efficiently lyse quiescent cells: 425-600 uM glass beads (Sigma, G8772) were added to samples in phenol-lysis buffer such that glass beads accounted for 1/3 of total sample volume. Greater than 50% of empty space was maintained in sample tubes to ensure efficient lysis. Samples were then vortexed for 1 min in 10 min intervals for 1 hour. rRNA depletion was performed using the Ribo-Zero (Yeast) rRNA Removal Kit (Illumina, San Diego, CA). Libraries were prepared with TruSeq Stranded Total RNA kit and purified using a Axygen AxyPrep MAG PCR Clean-Up Kit (Axygen). Illumina reads were mapped to the S288c genome substituted with SNPs from YPS1009 as called in Sardi et al.^154^, using bwa-meme^155^. Read counts for each gene were calculated by HT-Seq^156^. Normalization was conducted by setting the slope of *Sz. pombe* reads across samples to 1.0. Statistical analysis of log_2_(fold change) transcript abundances was done in edgeR^157^ taking genes with a false discovery rate (FDR)<0.05 as statistically significant. Genes shown in Fig 1E were defined as those significantly repressed (FDR<0.05) in the euploid strain and statistically significantly higher (FDR <0.05) in all four aneuploids, in at least one time point comparing that aneuploid to the euploid. Hierarchical clustering was performed using Cluster 3.0^158^ and visualized in Java Treeview^159^. Data represent the average of biological duplicate and are available in GEO Accession #GSE269236. Data for Figure 1D are available in Dataset 2.

### Genetic screen

Dual-marked YPS1009_euploid (AGY1611) and _Chr12 strains (AGY1612) were transformed with Moby 1.0 low-copy expression library^64^ and an aliquot removed as the starting pool. Cells were grown in biological triplicate for 28 days in YPD medium + G418 to maintain plasmids. A portion of each culture was harvested at 24 hours and 28 days after cultures completed 3 generations of growth. The harvested portions were plated on multiple plates of YPD + G418 to select for cells that were viable and able to form colonies, thus representing quiescent cells. After 48 hours of growth, lawns were scraped, collected, and flash frozen. Plasmid DNA was collected from the starting pools, day 1, and day 28 samples using Zymoprep Yeast Plasmid kit (Zymo Research, D2004), with the following changes: 425-600 uM glass beads were with the lysis reagent, and samples were vortexed for 10’’ three times during lysis. Samples were incubated on ice for 30 min after adding neutralization buffer. Barcodes were sequenced as previously described^160,161^. EdgeR was used to TMM normalize samples as previously described^157^. Genes with a significant positive log_2_(fold change) (FDR < 0.05) in barcode abundance at day 28 versus starting pool were considered beneficial. We then selected genes with a 2-fold or greater abundance difference between average YPS1009_Chr12 sample versus average euploid sample, which resulted in 59 candidate genes. Hierarchical clustering was performed on the log_2_(fold change) abundance differences using Cluster 3.0^158^ and visualized using Java TreeView^159^. Data are available in GEO Accession #GSE269237.

### Western blotting

Yeast strains were grown as described above, with the following additions: G418 was used to maintain the ribosome stalling reporter, and cells exposed to NTC were treated with 1 ug/mL NTC after cells reached mid-log phase. 2 OD units were harvested and flash frozen. Western blots were developed using anti-GFP (Abcam, ab290) for samples containing the ribosome stalling reporter, anti-ubiquitin (Milipore Sigma, MAB-1510), or anti-HA (Cell Signaling Technology, C29F4). Blots were developed on a Li-COR Odyssey instrument (Model 9120). Li-COR Odyssey software was used to quantify signal intensity of GFP and ubiquitin antibodies. Repeated attempts to blot against common loading controls were unsuccessful in quiescent cultures; therefore, Ponceau S signal was used to normalize protein loading levels, as performed by others^162^. Ponceau S signal was quantified using FIJI (ImageJ).

### Proteasomal activity

Proteasome-glo Cell-Based Assays (Promega) was used to measure proteasomal activities. Reagents were prepared according to manufacturer’s instructions. An equivalent number of yeast cells were flash frozen then lysed via vortexing with 500 uM glass beads on ice. Cell lysate and Proteasome-glo reagents were combined 1:1 in an opaque, white-walled 384-well plate (Corning). Luminescence was measured using a Tecan M1000 Pro.

### Ribosome profiling and analysis

Cells were harvested using vacuum filtration with Whatman Nylon filters (Cytiva, 7410-004). Cells were immediately scraped from filters, transferred to eppendorf tubes, and immediately flash frozen. Collection time was < 60 seconds. Ribosome profiling was performed by Ezra Biosciences as previously described^163^. Samples were sequenced on an Illumina Novaseq instrument and processed as described in Schuller et al.^164^ as follows: reads were trimmed with CutAdapt (version 3.5)^165^ with command j 8 -g ^GGG -a A{10} -n 2 -m 15 --max-n=0.1 --discard-casava. Reads with poor quality at the 5’ end base (quality score <= 10) were removed, reads were mapped to noncoding RNAs from Schuller et al.^164^, and remaining unmapped reads were mapped to the YPS1009 genome^23^ using bowtie2 (version 2.5.1)^166^. The 5’ position of each read was scored, and the P site taken to be at 12 nt into the read^163^. Reads matched well to the expected frame in all samples (see Supplemental Fig S3 for examples). For each gene, read starts were summed for each position from -72 of the gene ATG and + 60 of each stop codon in the YPS1009 genome. Genes without an annotated ATG were omitted from analysis, as were genes with introns. Read counts were summed for each codon, incremented by 1 pseudocount, and then normalized to the sum read counts (with appropriate pseudocounts) in each gene body, from 60 nt (20 codons) into the gene to 60 nt (20 codons) from the 3’ end as done previously^44^. Genes with at least 50 reads per gene body were retained for further analysis.

The correlation between transcript profiles shown in Fig 3E was taken as the uncentered Pearson correlation for each transcript as measured in euploid and aneuploid, paired by replicate (Fig 3E). Significant differences in ribosome peaks across replicate-paired aneuploid-euploid samples was calculated using Fisher’s Exact test with Benjamini-Hochberg multiple test correction^167^, by comparing read count in each sample at a given codon to gene-body read counts for that transcript (# reads at that codon, # reads in the gene body, for euploid versus aneuploid in each replicate separately). Peaks more abundant in the aneuploid were taken as those with FDR < 0.05 and for which the normalized ratio of read counts at that codon was greater in aneuploids; vice versa for peaks more abundant in the euploid. Motif analysis in Fig 3H was performed as follows: codons whose normalized read count differed between euploid and aneuploid samples (FDR < 0.05) were combined across replicates, and peaks more or less abundance in aneuploids versus euploids were partitioned. Ten amino acids flanking each peak site were retrieved from the YPS1009 proteasome. The frequency of each amino acid (and stop codon) at each position in the matrix was calculated as the number of occurrences of that amino acid divided by the number of peaks scored. Count and total values were compared at each position in the matrix for euploids versus aneuploids, using Fishers exact test and Benjamini-Hochberg FDR correction. Enrichments shown in Figure 3H were taken as the log2(fold difference) in frequency and shown only for statistically significant positions (FDR < 0.05, Fisher’s exact test). Quiescent aneuploids showed significant differences in amino acid composition at peaks detected in quiescence (Fig 3H); there were no significant differences for a comparable analysis done for log-phase cells (FDR > 0.05 in all cases). Data are available in GEO Accession #GSE269238.

## ACKNOWLEDGEMENTS

We thank Doug Kellogg for providing CLN3-6X-HA strain and members of the Gasch Lab for helpful discussions. This work was supported by NIH grant R01GM14975 to APG.

## Author Contributions

L.E.E and A.P.G. designed research; L.E.E, J.H., H.H., N.P performed research; L.E.E, M.P, and A.P.G analyzed data; L.E.E. and A.P.G. wrote the paper.

---

**Figure S1.**
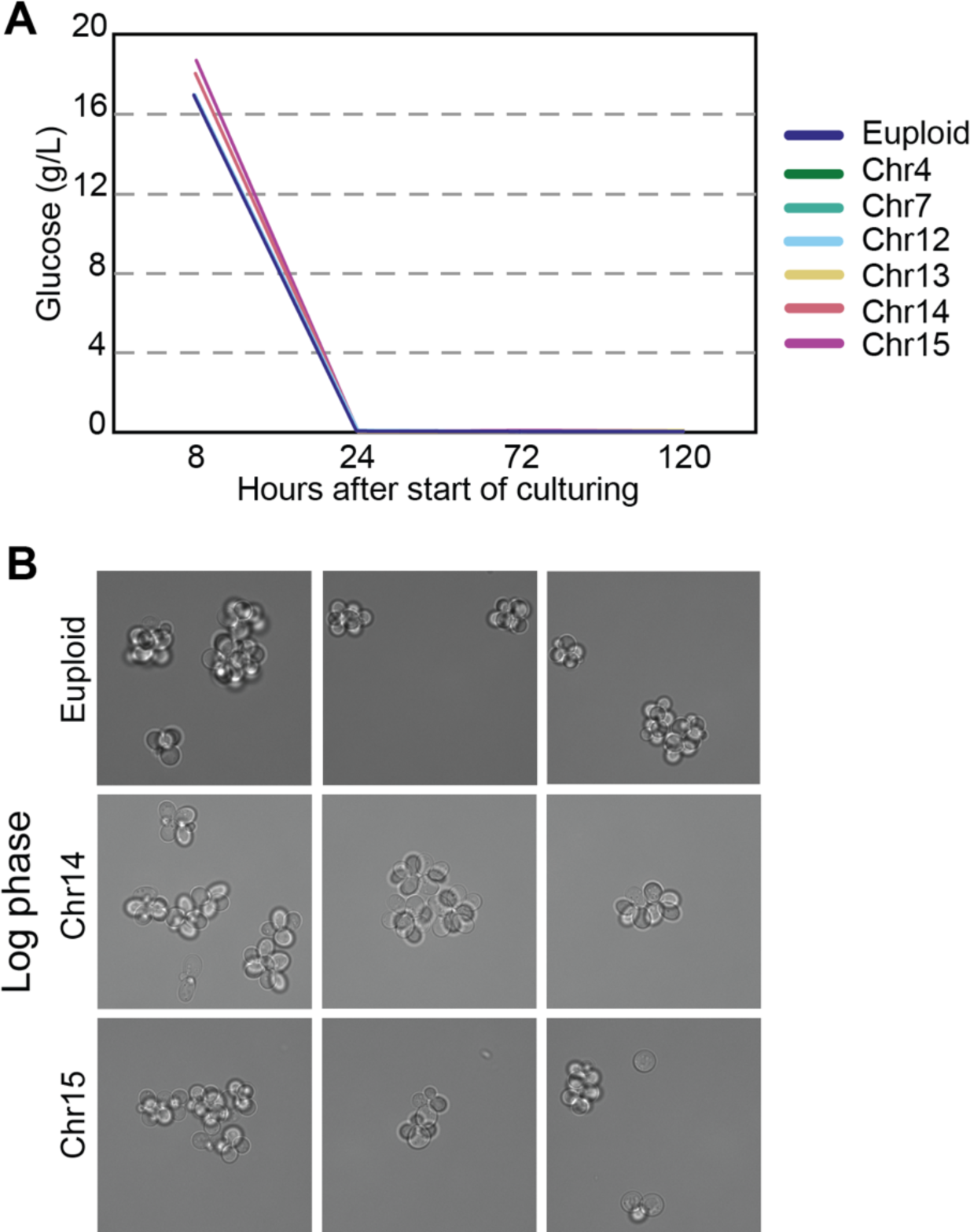
Aneuploid and euploid yeast cells grow similarly during proliferative growth. A) HPLC analysis of glucose concentration in euploid and aneuploid strains from 8 to 120 hours after start of culturing. Euploid, YPS1009_Chr12, Chr14, and Chr15 were measured at 8 hours and beyond; other aneuploids were measured at 24 hours and beyond. All cultures showed <0.04 g/L glucose at 24 hours; some curves are superimposable in the figure. B) Representative brightfield images of live euploid and aneuploid cells during log-phase demonstrate that aneuploids do not show unusual morphologies during log phase.

**Figure S2.**
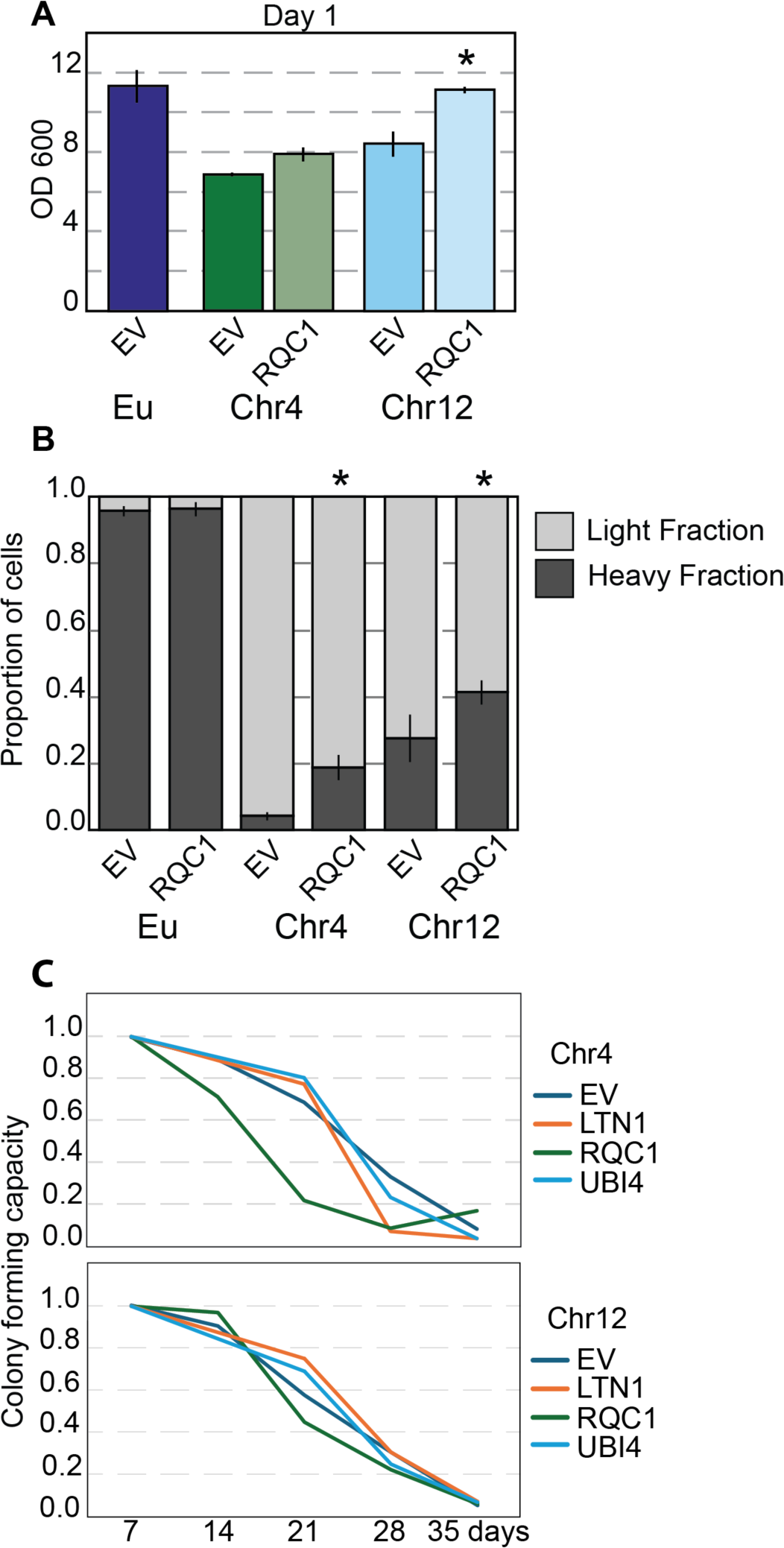
Overexpression of RQC plasmids bolsters some quiescence phenotypes in aneuploids. A) OD_600_ of euploid and aneuploid cultures harboring either empty vector (EV) or *RQC1* plasmid after 1 day of culturing. Asterisk, p<0.05, paired T-test, n = 3. B) Proportion of dense and light cells after 4 days. Asterisk, p<0.05, Chi-squared test, n = 2. C) Average fraction of colony forming units normalized to day 7 of Chr4 or Chr12 aneuploids harboring either empty vector or *LTN1, RQC1,* or *UBI4* plasmids.

**Figure S3.**
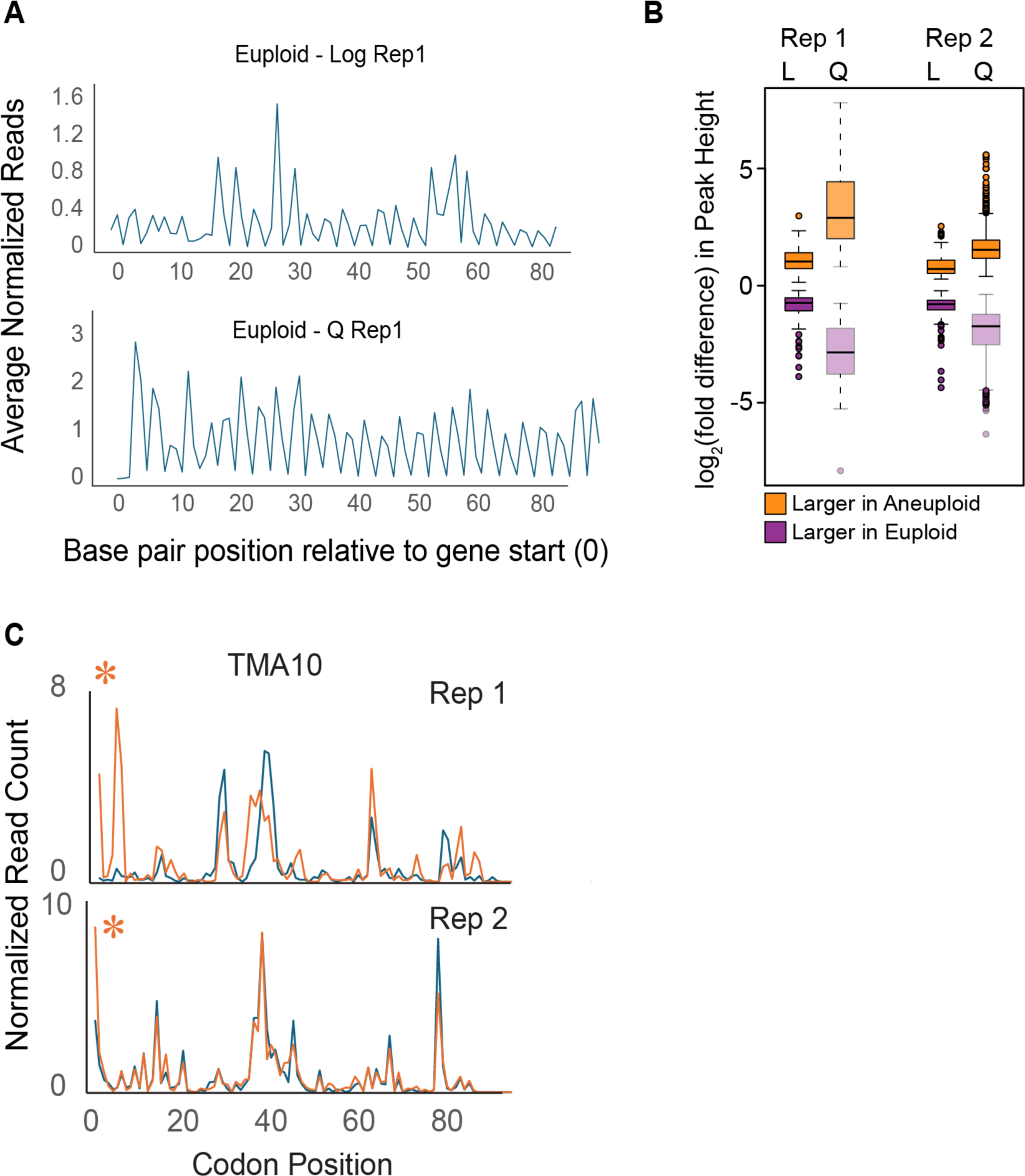
Ribosome profiling examples. A) Representative average read counts across all transcripts shows frame alignment across transcriptomes. B) Distribution of log_2_(fold difference) in normalized read counts (“Peak Heights”) for peaks scored as higher read count (normalized to gene body, see Methods) in aneuploids (orange) or in euploids (purple) in log phase (day 1) or quiescence (day 4) in two different replicates. In both replicates, a higher fraction of interrogated peaks were called significant during quiescence than log-phase and a higher fraction of those peaks had larger fold-differences in normalized read count. C) Representative traces at one transcript with significant aneuploid-enriched peak at the same position in both replicates (orange asterisk, FDR <0.05 in both replicates).

**Figure S4.**
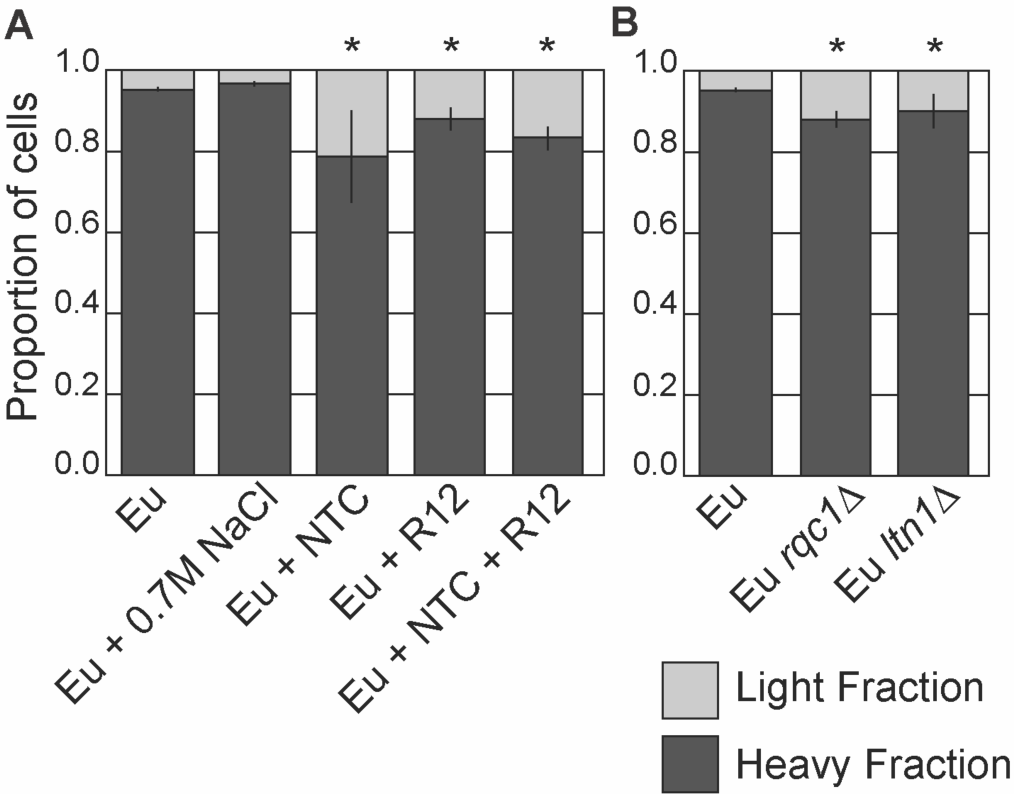
**Taxing RQC in euploid cells disrupts quiescence phenotypes**. A-B) Proportion of dense and light cells after 4 days. Asterisk, p<0.05, Chi-squared test, n = 2-4.

**Figure S5.**
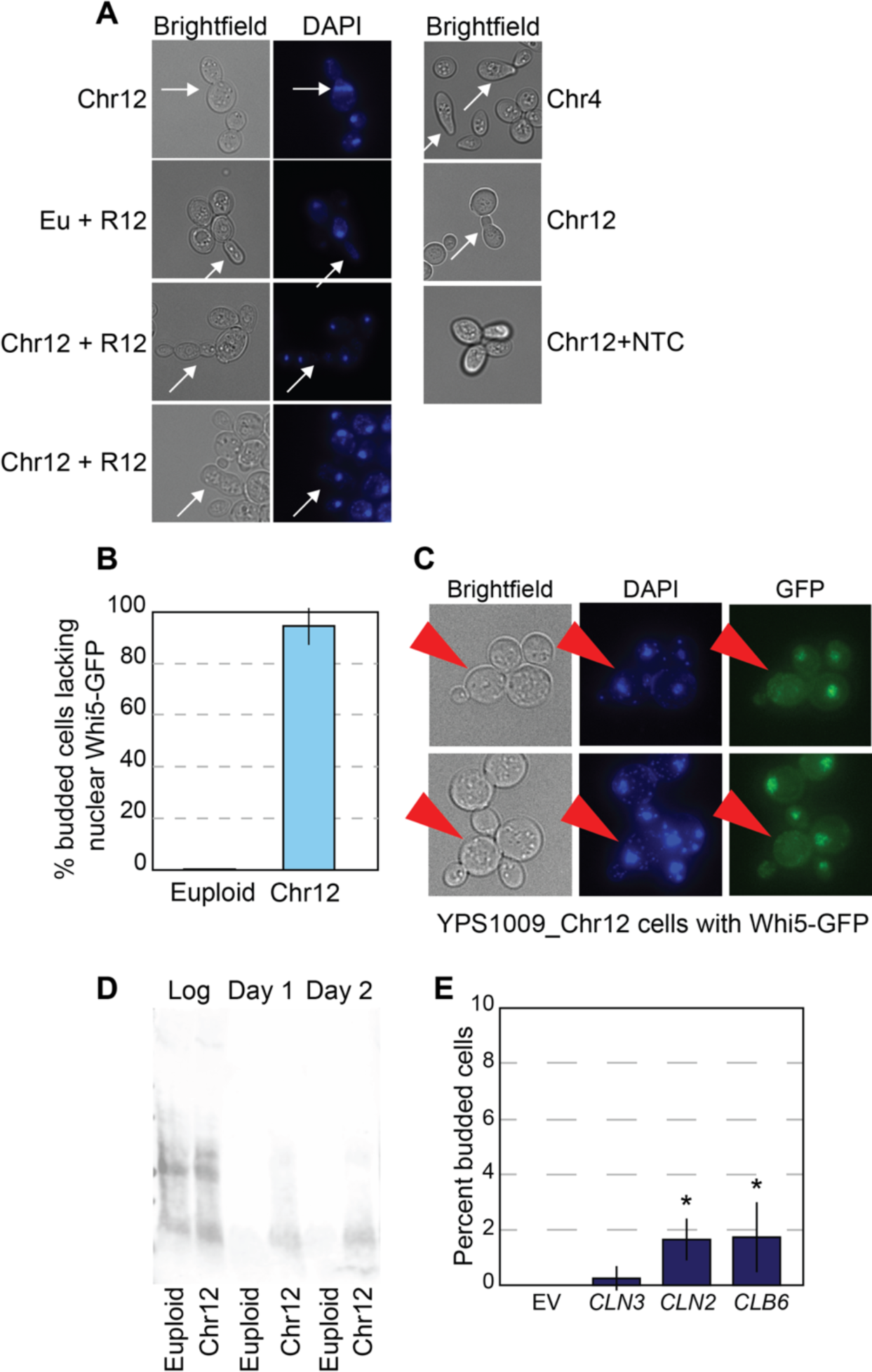
Cell cycle defects in aged aneuploid cells. A) Brightfield and DAPI images of euploid and aneuploid cells with notable morphology defects or nuclei polarity failures (white arrows). B) Percent of budding cells lacking proper nuclear localization of Whi5-GFP at 2 days (n = 3). C) Representative brightfield and fluorescent images of budding Chr12 cells that lack nuclear Whi5-GFP at 2 days (red arrows). D) Anti-HA western blot of Cln3-6xHA tagged euploid and YPS1009_Chr12 strains during log-phase, 1 and 2 days after start of culturing. E) Percent budding cells at 2 days in euploid cells harboring empty vector or *CLN3*, *CLN2*, or *CLB6* plasmids. Asterisk, p<0.05, unpaired T-test, n = 4-6.

**Figure S6.**
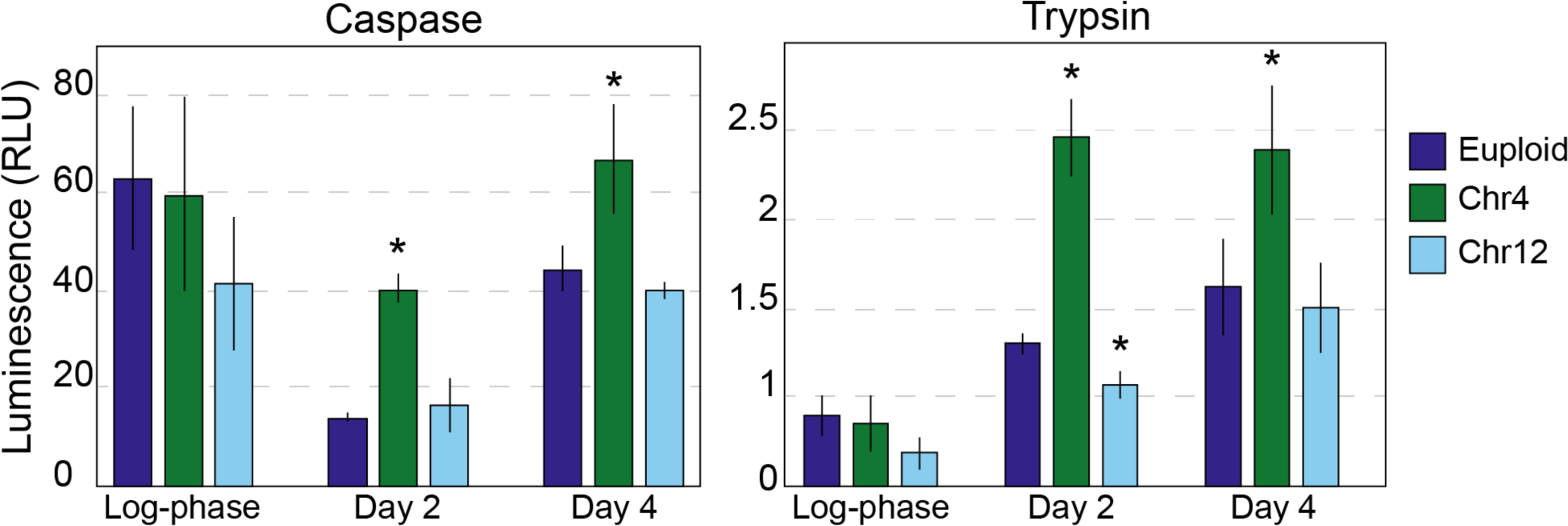
Euploid and aneuploid strains exhibit comparable *in vitro* proteasomal activities. *In vitro* caspase- and trypsin-like proteasomal activity in euploid and aneuploid strains at log-phase, and 2 and 4 days after start of culturing. Luminescence serves as an indicator of proteasomal cleavage of a luminescent substrate. Asterisk, p < 0.05 compared to euploid control, n=3; paired T-test.

**Figure S7.**
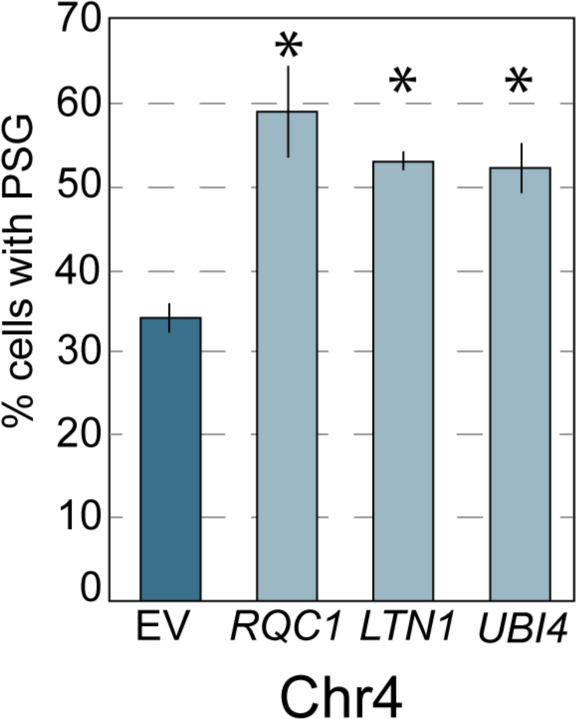
Percent of YPS1009_Chr4 cells with Pre6-GFP foci.

