## Supplemental-Figures for "Premature aging in aneuploid yeast is caused in part by aneuploidy-induced defects in Ribosome Quality Control"

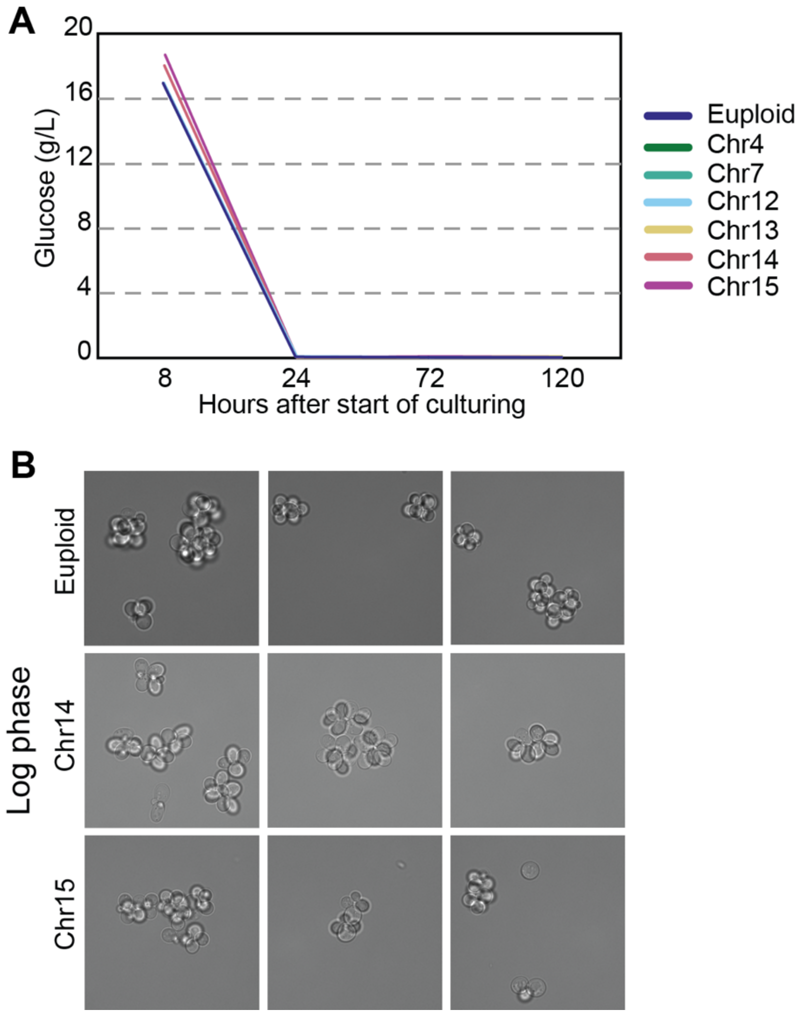


**Figure S1. Aneuploid and euploid yeast cells grow similarly during proliferative growth**. A) HPLC analysis of glucose concentration in euploid and aneuploid strains from 8 to 120 hours after start of culturing. Euploid, YPS1009_Chr12, Chr14, and Chr15 were measured at 8 hours and beyond; other aneuploids were measured at 24 hours and beyond. All cultures showed < 0.04 g/L glucose at 24 hours; some curves are superimposable in the figure. B) Representative brightfield images of live euploid and aneuploid cells during log-phase demonstrate that aneuploids do not show unusual morphologies during log phase.

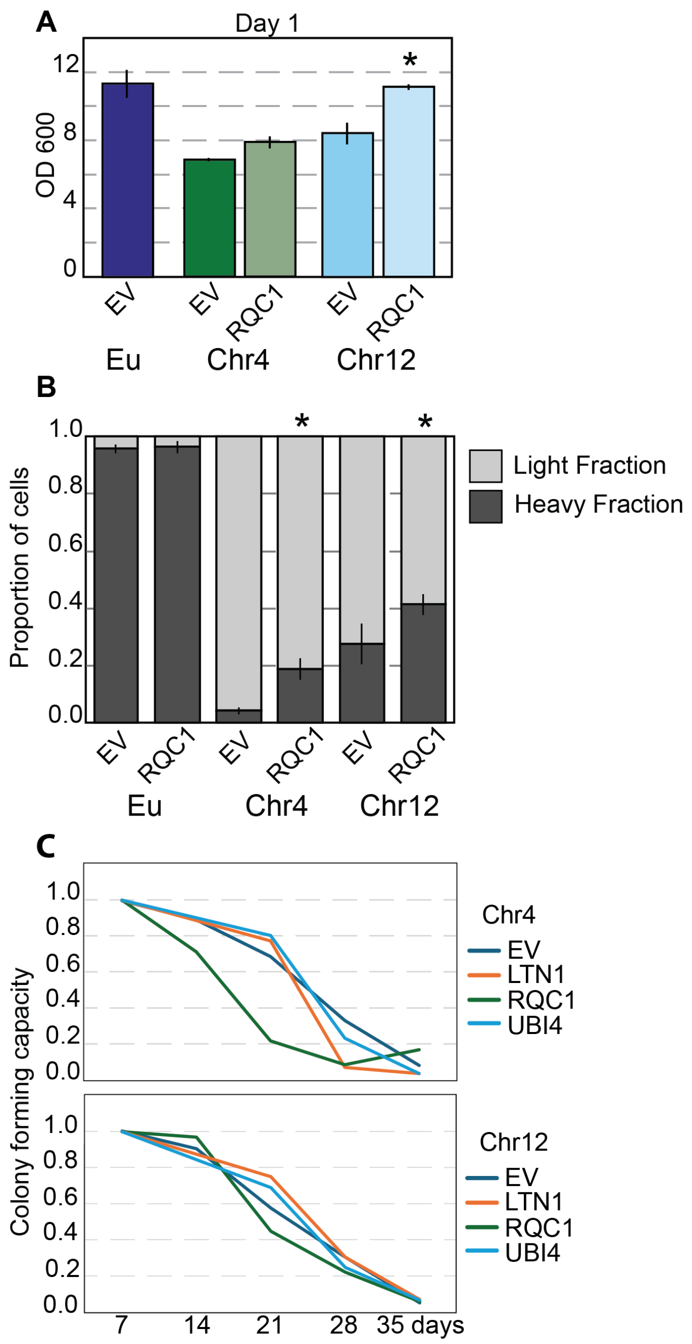


**Figure S2. Overexpression of RQC plasmids bolsters some quiescence phenotypes in aneuploids.** A) $OD_{600}$ of euploid and aneuploid cultures harboring either empty vector (EV) or *RQC1* plasmid after 1 day of culturing. Asterisk, $p<0.05$, paired T-test, n = 3. B) Proportion of dense and light cells after 4 days. Asterisk, $p<0.05$, Chi-squared test, n = 2. C) Average fraction of colony forming units normalized to day 7 of Chr4 or Chr12 aneuploids harboring either empty vector or *LTN1, RQC1,* or *UBI4* plasmids.

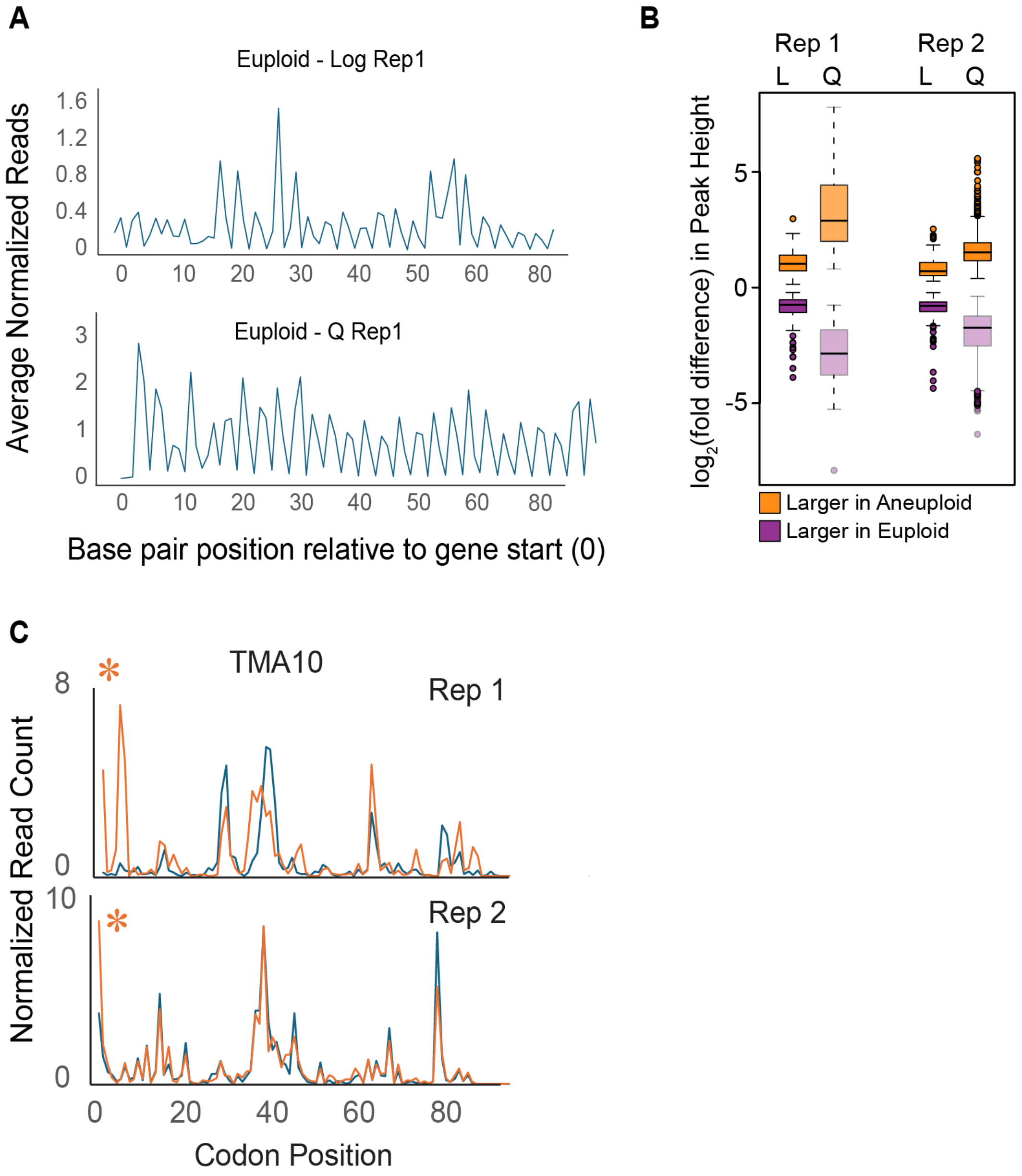


**Figure S3. Ribosome profiling examples.** A) Representative average read counts across all transcripts shows frame alignment across transcriptomes. B) Distribution of $\log_2$(fold difference) in normalized read counts (“Peak Heights”) for peaks scored as higher read count (normalized to gene body, see Methods) in aneuploids (orange) or in euploids (purple) in log phase (day 1) or quiescence (day 4) in two different replicates. In both replicates, a higher fraction of interrogated peaks were called significant during quiescence than log-phase and a higher fraction of those peaks had larger fold-differences in normalized read count. C) Representative traces at one transcript with significant aneuploid-enriched peak at the same position in both replicates (orange asterisk, FDR <0.05 in both replicates).

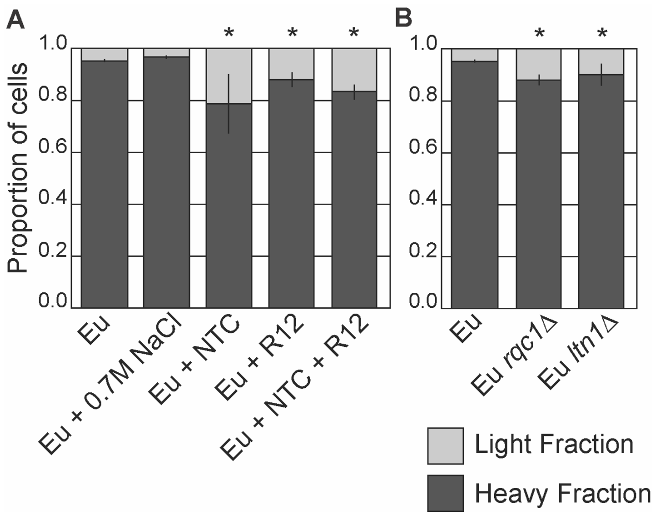


**Figure S4. Taxing RQC in euploid cells disrupts quiescence phenotypes**. A-B) Proportion of dense and light cells after 4 days. Asterisk, $p<0.05$, Chi-squared test, n = 2-4.

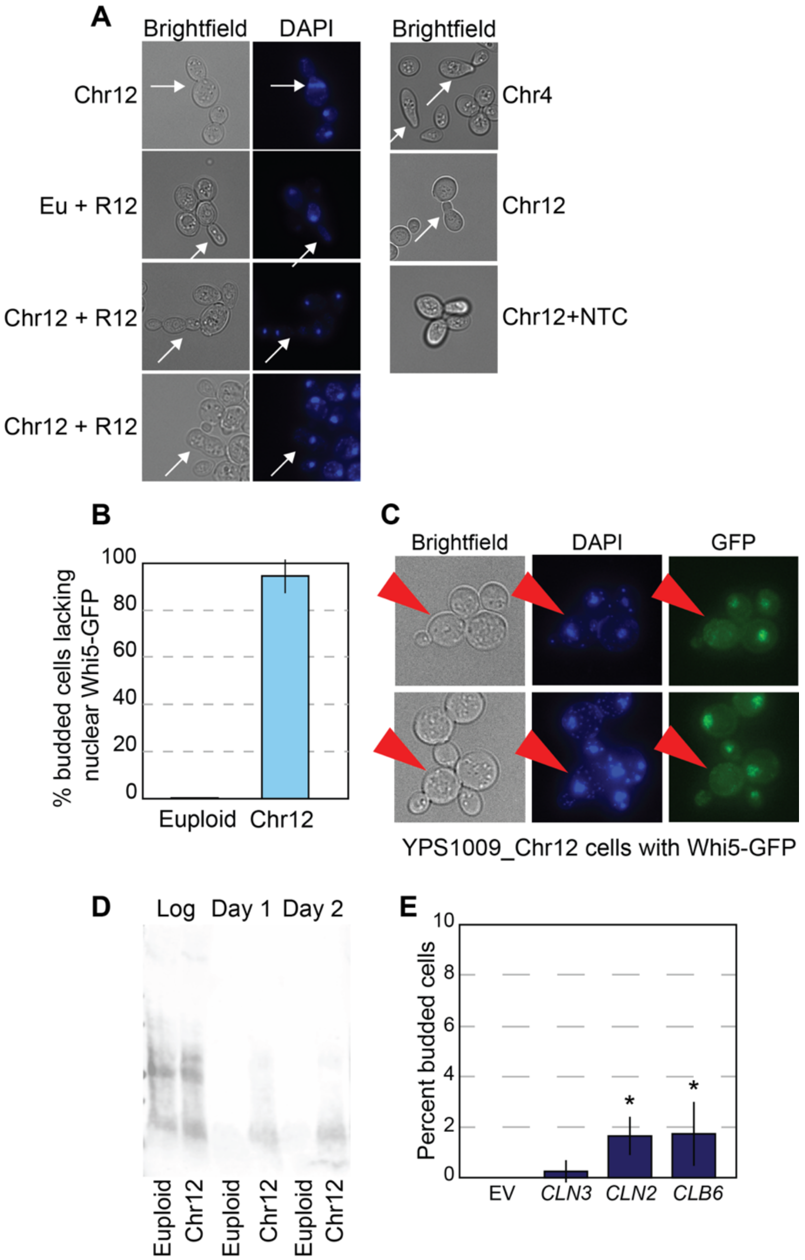


**Figure S5. Cell cycle defects in aged aneuploid cells.** A) Brightfield and DAPI images of euploid and aneuploid cells with notable morphology defects or nuclei polarity failures (white arrows). B) Percent of budding cells lacking proper nuclear localization of Whi5-GFP at 2 days (n = 3). C) Representative brightfield and fluorescent images of budding Chr12 cells that lack nuclear Whi5-GFP at 2 days (red arrows). D) Anti-HA western blot of Cln3-6xHA tagged euploid and YPS1009_Chr12 strains during log-phase, 1 and 2 days after start of culturing. E) Percent budding cells at 2 days in euploid cells harboring empty vector or *CLN3*, *CLN2*, or *CLB6* plasmids. Asterisk, $p<0.05$, unpaired T-test, n = 4-6.

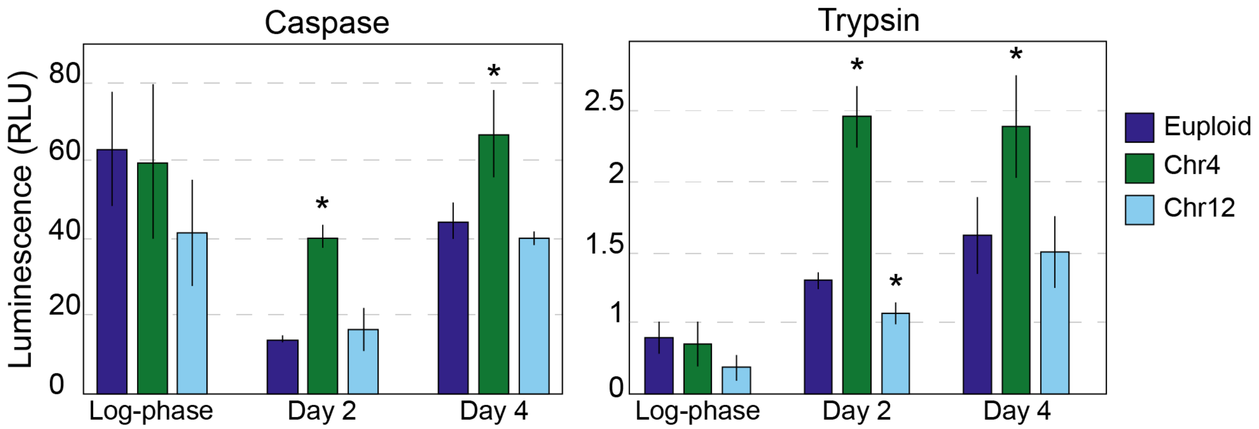


**Figure S6. Euploid and aneuploid strains exhibit comparable *in vitro* proteasomal activities.** *In vitro* caspase- and trypsin-like proteasomal activity in euploid and aneuploid strains at log-phase, and 2 and 4 days after start of culturing. Luminescence serves as an indicator of proteasomal cleavage of a luminescent substrate. Asterisk, $p < 0.05$ compared to euploid control, n=3; paired T-test.

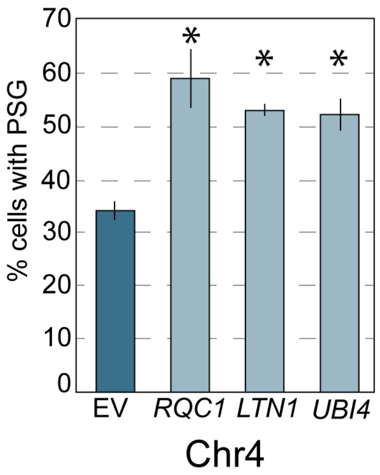


**Figure S7.** Percent of YPS1009_Chr4 cells with Pre6-GFP foci.
